# CD300LG is a receptor for triglyceride-rich lipoproteins that facilitates postprandial lipid clearance

**DOI:** 10.1101/2025.08.08.669356

**Authors:** Mitchell E. Granade, Kevin M. Tveter, Hye In Kim, Alexandra R. Yesian, Joseph Fowler, Youngwook Ahn, Jeffrey Culver, Kelly Tam Neale, Nalissa Amar, Michael Mazzocca, Alexis R. White, Danielle Archambault, Rajni Sonavane, Brigitte Laforest, Mary Piper, Rachel J. Roth Flach, Xin Rong

## Abstract

Circulating triglycerides are principally transported by triglyceride-rich lipoprotein particles (TRLs) including very-low-density-lipoproteins (VLDL) and chylomicrons and require the activity of lipoprotein lipase for appropriate lipid processing and cellular uptake. Despite known genetic links between *CD300LG* variants and altered lipid profiles, the functional role of CD300LG in lipid metabolism remains unclear. In this study, we identify CD300LG as a crucial receptor for TRLs. Human genetic analyses reveal that reduced CD300LG protein levels are causally linked with CAD risk and increased number, diameter, and TAG concentration of TRLs. In mice, CD300LG deficiency results in postprandial hypertriglyceridemia independent of changes in VLDL secretion, intestinal lipid absorption, or lipoprotein lipase activity. Mechanistically, CD300LG acts as a receptor for TRLs through a direct interaction with ApoA4 to facilitate TRL clearance at the microvascular endothelium. These findings elucidate new functions for both CD300LG and ApoA4 and advance our general understanding of triglyceride metabolism.

## Introduction

Despite the efficacy of current therapeutics targeting low-density lipoprotein (LDL) cholesterol levels in reducing coronary artery disease (CAD) risk, significant residual cardiovascular risk persists.^1^ Elevated circulating triglycerides (TGs) are associated with this residual cardiovascular risk, and postprandial TG levels show a stronger correlation with CAD risk than fasting TG levels.^2,3^ Therefore, understanding the underlying mechanisms involved in TG clearance will enable novel therapeutic discovery to reduce residual CAD risk.

CD300LG (CLM-9, NEPMUCIN, TREM-4) is a type 1 membrane protein that contains a small intracellular domain, a single transmembrane domain, a mucin-like domain, and an immunoglobulin domain. CD300LG is highly expressed within microvascular endothelial cells (ECs), but not in medium-sized or larger vessels.^4–7^ In ECs, CD300LG has been primarily characterized for its role as a ligand for L-selectin, which facilitates lymphocyte adhesion and trafficking at high endothelial venules.^4,5,8^ Recent genetic studies have identified a missense mutation (Arg82Cys) in the *CD300LG* gene that is associated with decreased fasting high-density lipoprotein cholesterol (HDL-c) and increased fasting TGs.^9–15^ Furthermore, genome-wide association studies (GWAS) demonstrated that SNPs at the *CD300LG* locus were associated with alterations in plasma lipid profiles.^10,16^ Although CD300LG has been implicated in lipid binding *in vitro* – specifically to phospholipids – its functional role in TG clearance remains undefined.^17^

In the present study, we performed comprehensive human genetics analyses using datasets with detailed lipoprotein profiles. Our analyses revealed that reduced CD300LG protein levels causally drive an increase in the number, size, and triglyceride content of triglyceride-rich lipoprotein particles (TRLs), contributing to elevated CAD risk. To validate these findings *in vivo*, we generated *Cd300lg*-deficient mice, which exhibited marked intolerance to triglyceride challenge and impaired postprandial lipid clearance independent of changes in VLDL secretion, intestinal lipid absorption, or lipoprotein lipase (LPL) expression and activity. Mechanistically, we demonstrated that CD300LG binds TRLs via a direct interaction involving the c-terminus of the apolipoprotein ApoA4. Collectively, our findings suggest CD300LG is a novel receptor for TRLs that facilitates postprandial lipid clearance in humans and mice.

## Results

### Lipoprotein profile in CD300LG R82C carriers suggests that CD300LG loss-of-function impairs TRL metabolism

Previous GWAS studies reported an association between the Arg82Cys (R82C) variant in *CD300LG* and plasma TG and HDL-c levels, suggesting a role of CD300LG in the regulation of plasma lipids.^10,18,19^ Using the large-scale proteomics GWAS in UK Biobank,^20^ 8 independent signals near *CD300LG* (cis-pQTLs) were identified to be associated with circulating CD300LG protein levels upon fine-mapping (Fig. S1A). This included the R82C mutation, which was associated with reduced CD300LG protein levels, suggesting that CD300LG loss-of-function is associated with increased TG levels and reduced HDL-c levels (Fig. 1A). Several other cis-pQTLs were also colocalized with plasma lipid signals with consistent directionality (Fig. S1B). Mendelian randomization analyses using the cis-pQTLs further confirmed the causality and directionality of the effect of CD300LG on plasma lipids (Fig. S1C-D). To better understand how CD300LG influences plasma lipid levels, the association of R82C with detailed lipoprotein profiles measured by NMR was examined in UK Biobank.^21,22^ R82C is associated with an increased number of VLDL particles and increased VLDL diameter, but decreased number of HDL particles and decreased LDL and HDL diameters (Fig. 1A,B, Table S1). While total cholesterol (TC) levels are lower in the R82C carriers, both TG and cholesterol levels in VLDL particles are elevated (Fig. 1C-D, Table S2). Overall, the lipoprotein profile in R82C carriers suggests that CD300LG may specifically alter TRL metabolism and turnover.

**Figure 1.**
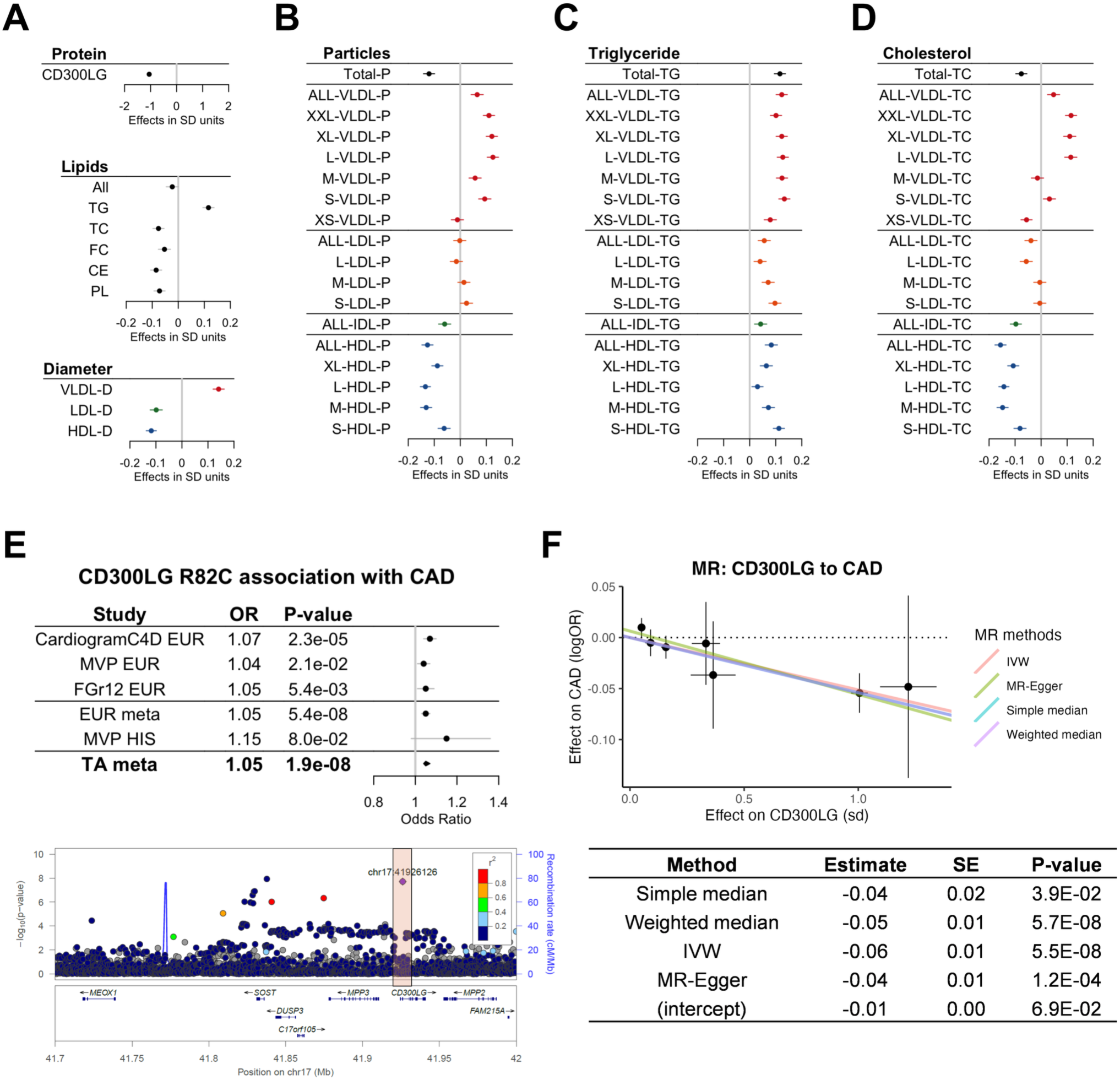
Human genetic associations at the CD300LG locus suggest a role in TRL metabolism and CAD risk. A) Effect of R82C variant on CD300LG protein, lipids (TGs, total cholesterol (TC), free cholesterol (FC), cholesterol esters (CE), and phospholipids (PL)), and lipoprotein particle diameter. B-D) Effect of R82C on lipoprotein particle concentration (B), TG (C), and cholesterol (D) across lipoprotein fractions. E) Meta-analysis of R82C association with CAD across three studies. F) Mendelian randomization analysis using CD300LG cis-pQTLs as the exposure and CAD risk as the outcome. Error bars in forest plots represent 95% confidence intervals. The dots and error bars in Mendelian randomization plot represent the effect size and standard error of the association between the variants and the exposure and outcome traits.

### Mendelian randomization analysis suggests a causal effect of CD300LG on CAD risk reduction

To investigate the potential effect of CD300LG on CAD through its effect on TRLs, we meta-analyzed the association of R82C with CAD from prior GWAS studies (CARDIoGRAMplusC4D^23^, Million Veteran Program (MVP)^24^, FinnGen (FGr12)^25^. Upon meta-analysis, R82C was significantly associated with a 5% higher risk of CAD (Fig. 1E, Table S3). Additionally, we performed Mendelian randomization analysis using the CD300LG cis-pQTLs as instrumental variables of the exposure and CAD as the outcome. This further confirmed the causal effect of CD300LG on CAD, with higher CD300LG protein levels leading to decreased CAD risk (Fig. 1F, Table S2).

### Cd300lg knockout mice exhibit impaired postprandial lipid tolerance

To further explore the function of CD300LG in lipid metabolism *in vivo*, we generated a germline *Cd300lg* knockout mouse model (Cd300lg-KO) (Fig. S2A). Expression analysis confirmed the absence of *Cd300lg* mRNA across multiple tissues, including heart, epididymal adipose (eWAT), and skeletal muscle (gastrocnemius) (Fig. S2B). CD300LG protein expression was detected on ECs in WT mice but absent in Cd300lg-KO mice (Fig. S2C-D).

WT and Cd300lg-KO mice were fed either a chow or high-fat diet (HFD) for 18 weeks (Fig. 2A, G). Male Cd300lg-KO mice showed no significant differences from WT mice in body weight or food intake regardless of diet (Fig. 2B-C, H-I). No differences in plasma TGs or TC were observed on chow diet, and a modest increase in plasma TGs was detected in HFD-fed Cd300lg-KO mice (Fig. 2D-E, J-K). However, upon an oral fat tolerance test (OFTT), Cd300lg-KO mice on both chow and HFD had substantially higher peak plasma TG levels and delayed TG clearance compared with WT (Fig. 2F,L). To specifically investigate postprandial lipid metabolism, plasma TG levels were measured after an overnight fast followed by a 2-hour refeed. Chow- refed Cd300lg-KO mice showed no differences compared to WT; however, HFD-refed Cd300lg-KO mice exhibited dramatically increased postprandial plasma TGs with an obvious lipemic appearance to the plasma (Fig. 2M-O). Female Cd300lg-KO mice had a similar phenotype as male mice (Fig. S3). Notably, despite substantial impairment in postprandial lipid metabolism, there were no significant changes in fat and lean mass distributions (Fig. S4A-B, H-I), tissue weights for epididymal adipose (eWAT), liver, and heart (Fig. S4C-E, J-L), or glucose or insulin tolerance (Fig. S4F-G, M-N). Together, our findings demonstrate that CD300LG plays a highly specific role in regulating postprandial TG metabolism, especially after a meal with high lipid content.

**Figure 2.**
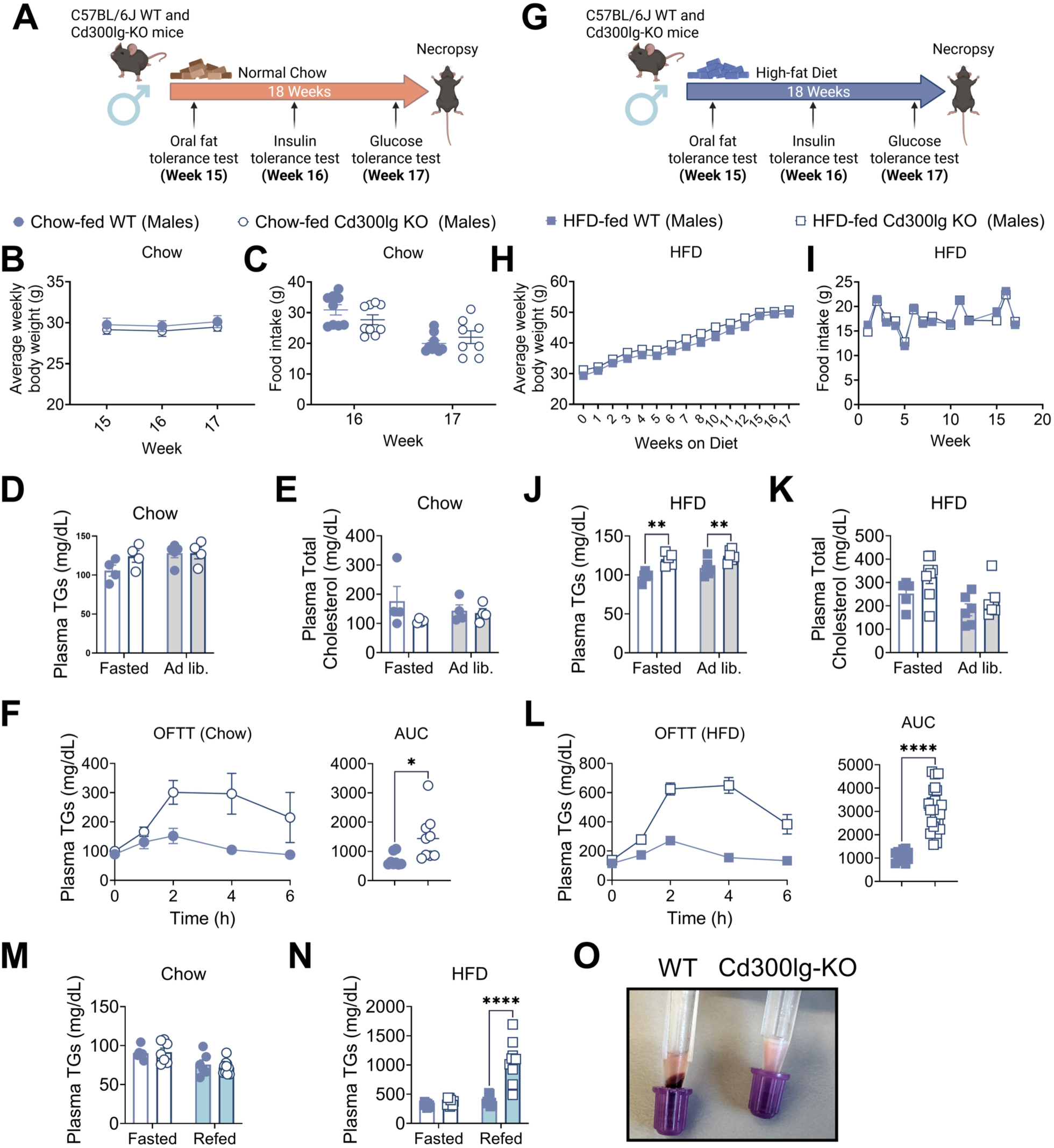
Cd300lg-KO mice exhibit postprandial lipid intolerance. A) Experimental design and timeline for WT and Cd300lg-KO male mice fed chow for 18 weeks post-weaning. B-C) Body weight (B) and food intake (C) for mice on chow at weeks 15-17. (N = 9). D-E) Plasma TGs (D) and total cholesterol (E) in fasted (white bars) and ad libitum (gray bars) chow-fed WT and Cd300lg- KO mice. F) Oral fat tolerance test (OFTT) in chow-fed mice; plasma TGs were measured at the indicated time points following olive oil gavage in overnight-fasted mice. G) Experimental design and timeline for WT and Cd300lg-KO male mice fed a high-fat diet (HFD) for 18 weeks post-weaning. H-I) Body weight (H) and food intake (I) for mice on HFD from weeks 0 to week 17 (N=9- 10). J-K) Plasma TGs (J) and total cholesterol (K) in fasted (white bars) and ad libitum (gray bars) HFD-fed WT and Cd300lg-KO mice. L) Oral fat tolerance test (OFTT) in HFD-fed mice, plasma TGs were measured at the indicated time points following olive oil gavage in overnight fasted mice. M) Plasma TGs in chow-fed mice fasted overnight (white bars) or following a 2-hour refeed on chow (light blue bars). N) Plasma TGs in HFD-fed mice fasted overnight (white bars) or following a 2-hour refeed on HFD (light blue bars). O) Representative image of plasma taken from WT and Cd300lg-KO male mice following a 2-hour refeed on HFD. Error bars denote S.E.M. *p<0.05, **p<0.01, ****p<0.0001 by t-test or two-way ANOVA as appropriate.

### Cd300lg does not impact VLDL secretion or intestinal lipid absorption

To better understand the mechanism underlying the impaired postprandial lipid metabolism in Cd300lg-KO mice, we first assessed the potential effects on VLDL production and intestinal lipid absorption. Lipoprotein lipase (LPL) mediates the hydrolysis of TGs in TRLs and is essential for their clearance.^26,27^ By inhibiting LPL with the lipase inhibitor, Poloxamer-407 (P-407), we assessed VLDL secretion and intestinal lipid absorption rates (Fig.3A).^28^ Cd300lg-KO mice showed no significant differences in VLDL secretion (Fig. 3B). Intestinal lipid absorption was assessed by administration of an oral lipid gavage after P-407 administration (Fig. 3A). Inhibition of LPL normalized postprandial plasma TG levels between Cd300lg-KO mice and WT mice (Fig. 3B), suggesting that the elevated postprandial TG levels in Cd300lg-KO mice are not due to altered intestinal lipid absorption or chylomicron (CM) production, but potentially the catabolism of CMs.

**Figure 3.**
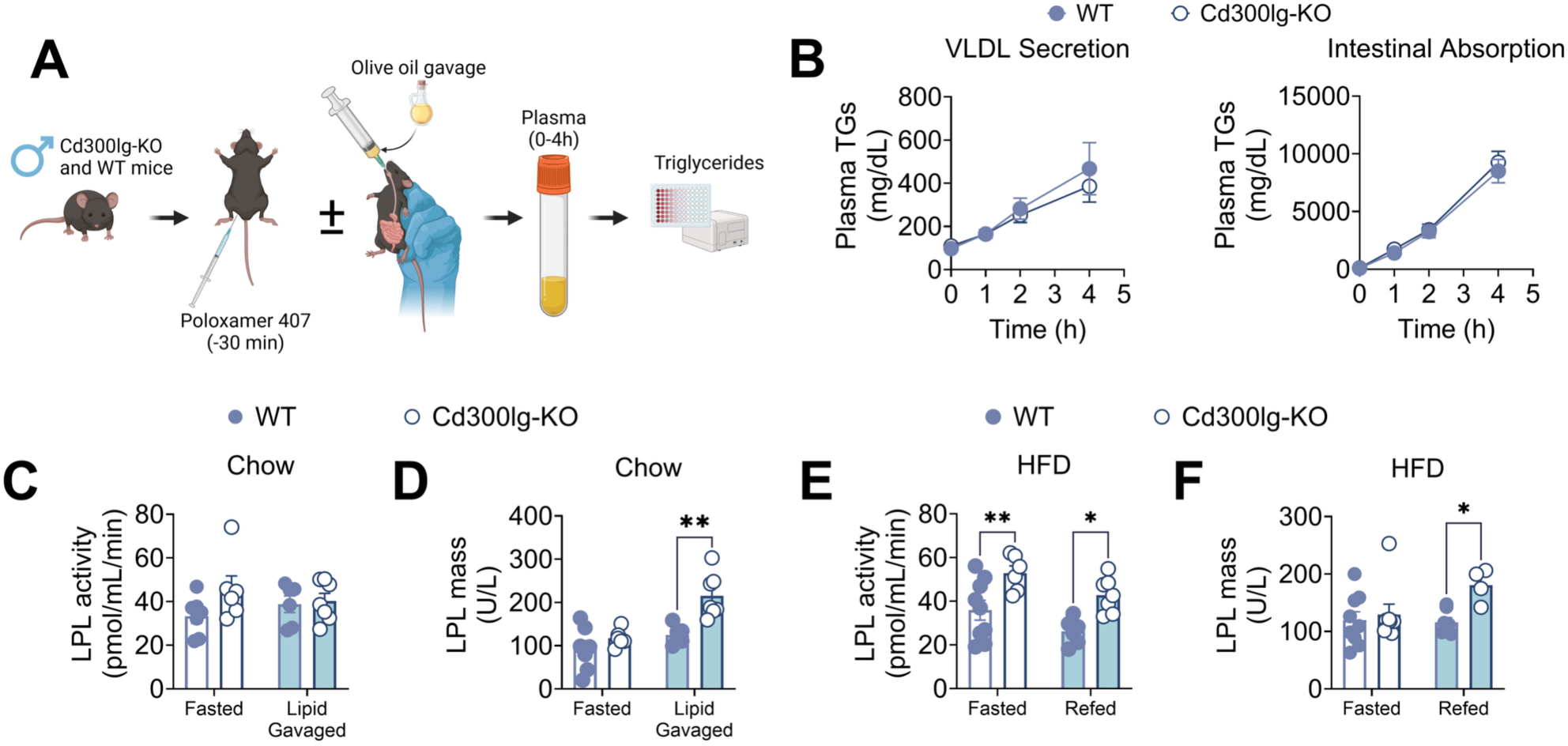
Cd300lg deficiency does not impair VLDL secretion nor intestinal lipid absorption and modestly increases LPL activity. A) Schematic overview of VLDL secretion and intestinal absorption assays. B) Plasma TGs in male WT (closed circles) and Cd300lg-KO (open circles) at baseline, 1, 2, and 4 hours post poloxamer 407 administration alone (VLDL secretion, N=6,7) or followed 30 minutes later by an oral gavage of olive oil (intestinal absorption, N=9,11). C-D) Post-heparin plasma LPL activity (C) or LPL protein levels (D) in chow-fed mice fasted overnight or 2-hours after an olive oil gavage. E-F) Post-heparin plasma LPL activity (E) or LPL protein levels (F) in HFD-fed mice fasted overnight or following a 2-hour refeed on HFD. Error bars denote S.E.M. *p<0.05, **p<0.01 by two-way ANOVA.

### Cd300lg expression correlates with LPL-related genes but does not impact LPL levels or activity

We next explored whether CD300LG functions in proximity to LPL. LPL is highly expressed in adipose, heart, skeletal muscle, and other tissues where significant lipid uptake occurs.^29^ A gene co-expression correlation analysis revealed a strong correlation between *CD300LG* and *GPIHBP1* (Fig. S5A). GPIHBP1 is responsible for the translocation of LPL across microvascular ECs to the apical membrane.^30–32^ The bulk tissue expression pattern of *CD300LG* was also highly similar to *GPIHBP1* and *LPL* (Fig. S5B). Further, single- cell RNAseq data from skeletal muscle and adipose demonstrated that the highest expression for both *CD300LG* and *GPIHBP1* is found within microvascular ECs, where LPL-mediated metabolism of TRLs occurs (Fig. S5C-D). We therefore hypothesized that CD300LG may be involved in the processing of TRLs by LPL.

Defects in either LPL activity or its translocation to the luminal surface of endothelial cells result in severe hypertriglyceridemia due to reduced clearance of TRLs.^26,27,30,33,34^ To determine whether Cd300lg-KO impacted luminal protein levels of LPL or its enzymatic activity, heparin was administered to both WT and Cd300lg-KO male mice to release LPL bound to ECs in the capillary lumen, and plasma LPL activity and protein levels were assessed. Chow-fed Cd300lg-KO mice showed no reductions in LPL activity or protein levels (Fig. 3C-D). Surprisingly, HFD-fed Cd300lg-KO mice exhibited moderately increased LPL activity and protein levels (Fig. 3E-F), suggesting the reduced postprandial TG metabolism in Cd300lg-KO mice is not due to defects in LPL presentation or activity. Furthermore, no differences were observed in mRNA levels of known LPL regulators except *Angptl8,* which was reduced in both the liver and eWAT of chow-fed Cd300lg- KO mice in the fasted state (Fig S6A-E). Finally, isolated TRLs (d<1.006) from WT and Cd300lg-KO mice were compared via proteomics, and no significantly differentially regulated proteins were observed (Fig. S6F, Table S4). Together, our data suggest that CD300LG does not regulate TRL metabolism through direct modulation of the presentation or activity of LPL at the vascular endothelium; instead, Cd300lg-KO mice exposed to chronic high-fat feeding exhibit increased LPL activity, perhaps as a compensatory response.

### CD300LG is sufficient for TRL binding to ECs

As Cd300lg-KO mice did not have defects in LPL expression or enzymatic activity, we next assessed whether CD300LG may be involved in the binding of TRLs to ECs. TRL binding to ECs has been previously ascribed to GPIHBP1-bound LPL; however, we hypothesized that CD300LG may play an additional, uncharacterized role in this process.^35,36^ Notably, immortalized EC lines do not express this machinery (neither GPIHBP1 nor LPL), and, similarly, *CD300LG* mRNA expression is also not detected in cultured ECs.^5,37^ Therefore, adenovirus was used to express either empty vector or human-CD300LG (hCD300LG) in hTERT-immortalized endothelial (TIME) cells. Dil-labeled human-derived isolated CMs were incubated with the TIME cells, and CM binding was measured with high-content imaging (Fig. 4A-B). Binding of Dil-CMs to TIME cells expressing empty vector was below the limit of detection; however, significant levels of CMs bound to cells expressing hCD300LG. To test whether this binding activity was exclusive to TRLs, we also tested the binding of Dil-labeled VLDL, LDL and HDL. While both VLDL and LDL bound to TIME cells expressing empty vector, expression of hCD300LG only increased the binding of VLDL, and HDL binding was not detected irrespective of hCD300LG expression (Fig. 4C). Thus, CD300LG expression is indeed sufficient to enable the binding of TRLs to ECs independent of LPL and GPIHBP1.

**Figure 4.**
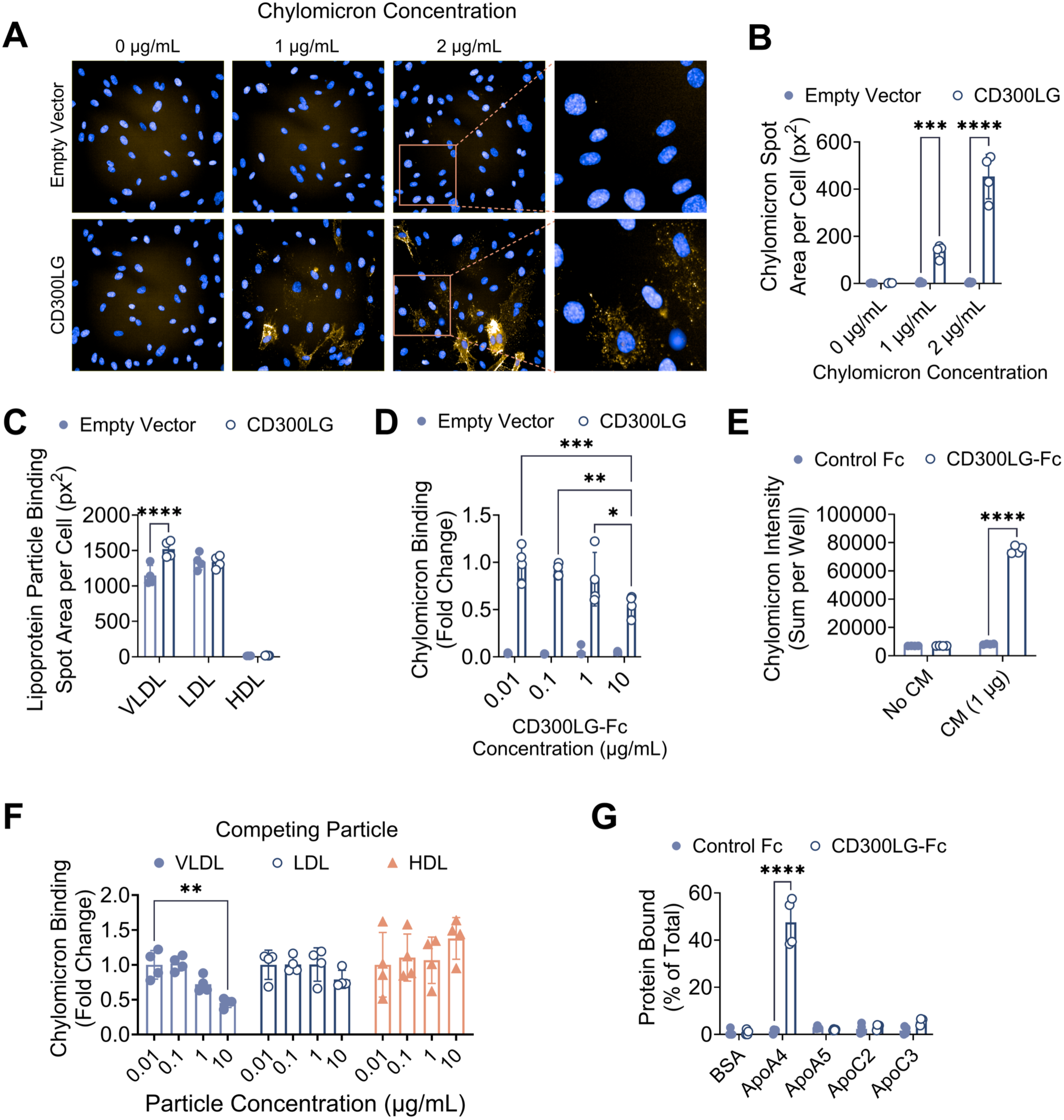
CD300LG directly binds TRLs and ApoA4. A-B) Representative images (A) and quantification (B) from high-content fluorescence microscopy of Dil-labeled chylomicrons (Dil-CMs) binding to TIME cells transfected with human CD300LG or empty vector. C) Quantification of Dil-labeled VLDL, LDL, and HDL (human) binding to human CD300LG or empty vector- transfected TIME cells by high-content imaging. D) Dose-dependent competition of DiI-CM binding to CD300LG by recombinant human CD300LG-Fc quantified by high-content imaging. E) Dil-CMs directly bind recombinant human CD300LG-Fc, but not control-Fc, immobilized on ELISA plates. Total fluorescence intensity was measured across entire wells. F) Binding of Dil-CMs following a pre-incubation of TIME cells overexpressing CD300LG with the indicated concentrations of unlabeled VLDL, LDL, or HDL. Binding was quantified by high-content imaging and normalized to cells incubated with Dil-CMs alone. G) Binding of hCD300LG-Fc or control Fc to purified apolipoproteins immobilized to ELISA plates. Error bars denote S.D. *p<0.05, **p<0.01, ***p<0.001, ****p<0.0001 by two-way ANOVA. All data are representative of a minimum of two independent experiments.

### CD300LG directly binds to chylomicrons and ApoA4

We next determined whether CD300LG was directly interacting with CMs. A recombinant soluble human CD300LG-Fc (hCD300LG-Fc) fusion protein was able to compete with the binding of Dil-CMs to TIME cells expressing CD300LG, suggesting that the CD300LG ecto-domain may be directly involved in this interaction (Fig. 4D). To further confirm the direct interaction, ELISA plates were prepared using either recombinant hCD300LG-Fc or an isotype control Fc as the capturing component. Substantial binding of Dil- CMs was detected in wells containing hCD300LG-Fc, but not the wells containing control Fc (Fig. 4E). These data suggested that CD300LG can indeed serve as a direct receptor for CMs.

The TIME cell binding data suggested that CD300LG only bound to CMs and VLDL. To confirm this specificity for TRLs, binding experiments between TIME cells and Dil-CMs were repeated in the presence of unlabeled VLDL, LDL, and HDL. Only unlabeled VLDL was able to competitively inhibit binding to Dil-CMs, while unlabeled LDL and HDL had no impact on Dil-CM binding (Fig. 4F).

Although CD300 family members have been previously shown to bind to lipids, CD300LG does not show selectivity for specific lipid species and thus selective binding to TRLs cannot be explained by the affinity to lipids.^17^ We therefore hypothesized that CD300LG interacted directly with CMs via a protein ligand, and thus investigated apolipoproteins specifically associated with TRLs. We focused on 4 prominent apolipoproteins with strong associations to CMs and VLDL that also have known roles in LPL regulation (ApoC2, ApoC3, ApoA5) or in CM production (ApoA4). ELISA-based screening identified specific binding between CD300LG and ApoA4 with no detectable binding to the other tested apolipoproteins (Fig. 4G). Together, these data suggest that CD300LG can directly interact with CMs, which may be mediated through a direct binding interaction with ApoA4.

### The C-terminus of ApoA4 is partially responsible for the binding activity with CD300LG

ApoA4, an exchangeable apolipoprotein produced as part of nascent CMs, plays a key role in CM production and metabolism.^38–40^ Structurally, ApoA4 contains a unique EQXQ repeat within its c-terminal region, which is highly conserved across species and absent from other apolipoproteins, such as ApoA5 (Fig. 5A). This domain was predicted as a potential protein-protein binding domain, but no definitive role has been ascribed.^41^ We hypothesized that this unique 16-amino acid repeating motif mediates the interaction between ApoA4 and CD300LG.

**Figure 5.**
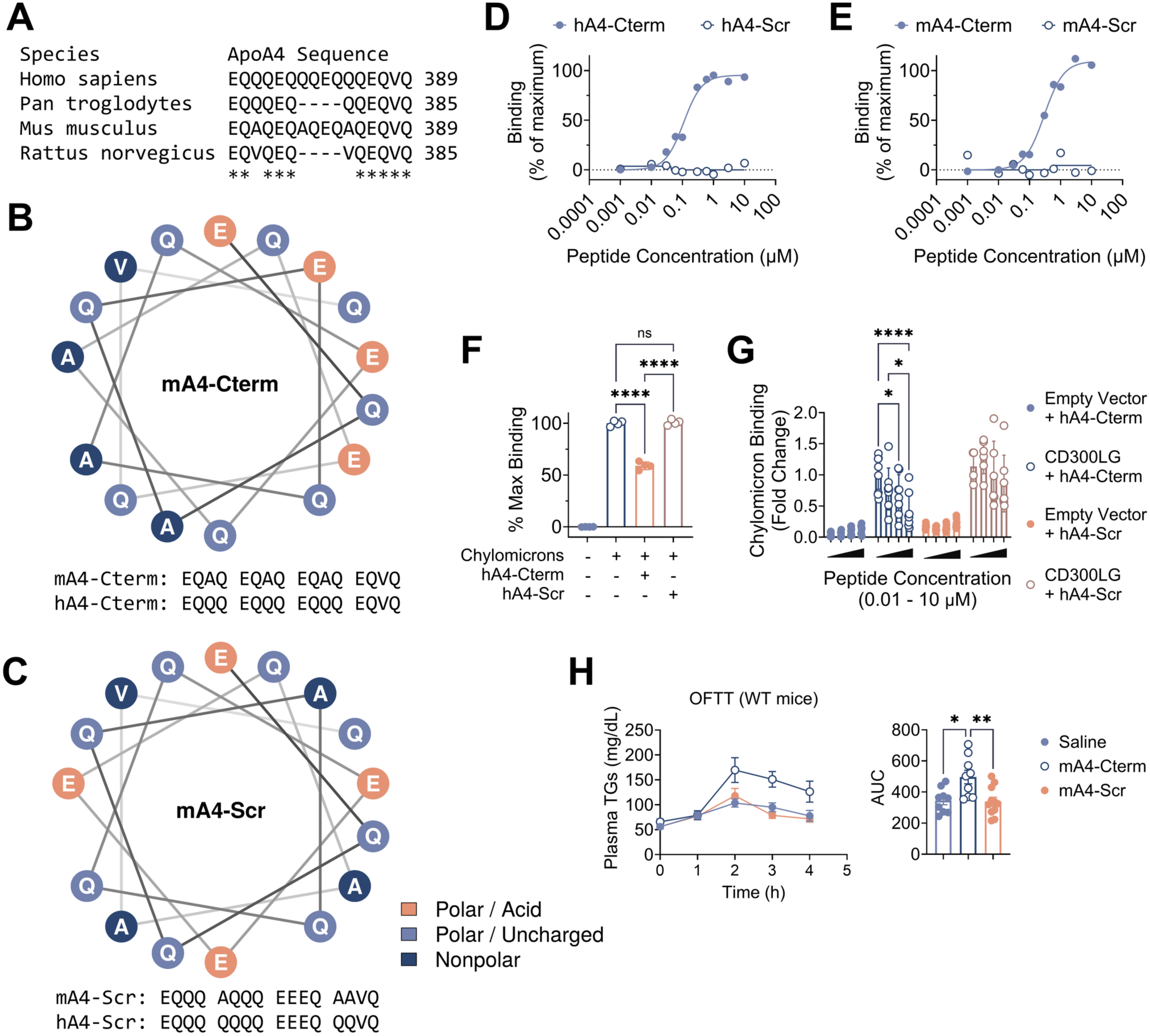
The c-terminus of ApoA4 contributes to CD300LG binding to TRLs. A) Sequence alignment of the conserved EQXQ motif in the C-terminal region of ApoA4 across species. B) Helical wheel diagram of the c-terminus of ApoA4 showing the one- sided charge distribution of glutamate residues created by the repeating EQXQ motif. C) Helical wheel diagram showing the redistribution of the charged glutamate residues within the “scrambled” control peptides. D-E) Direct binding of peptides corresponding to the human (D) or mouse (E) ApoA4 C-terminus (hA4-Cterm, mA4-Cterm) or scrambled controls (hA4-Scr, mA4- Scr) to recombinant human CD300LG-Fc. F) hA4-Cterm peptide inhibits the binding between human CD300LG-Fc and Dil-CMs. CD300LG-Fc coated plates were pre-incubated with peptides for 2 hours prior to the addition of CMs. G) hA4-Cterm peptide dose-dependently inhibits the binding of Dil-CM to CD300LG expressing TIME cells. TIME cells were pre-incubated with peptides for 30 minutes on ice prior to the addition of Dil-CMs. H) Oral fat tolerance test (OFTT) for WT mice treated with saline or 40 µg of mA4-Cterm or mA4-Scr 15 minutes before an oral gavage of olive oil (N = 9-11). AUC is quantified on the right. *p<0.05, **p<0.01, ****p<0.0001 by one-way or two-way ANOVA as appropriate. All data are representative of a minimum of two independent experiments. Data points in panels D and E represent the average of 2 independent experiments.

Synthetic peptides representing the mouse (mA4-Cterm) and human (hA4-Cterm) ApoA4 c-terminal motifs were generated. Both peptides are predicted to form a unique alpha-helix, with a highly uneven distribution of the negatively charged glutamate residues along one side of the helix (Fig 5B). To evaluate whether this negatively charged motif is required for the binding to CD300LG, we made specific changes to the sequence to disrupt the charge distribution while maintaining the amino acid composition to generate mouse (mA4-Scr), and human (hA4-Scr) “scrambled” controls (Fig. 5C). Both mouse and human peptides exhibited significant dose-dependent and saturable binding to hCD300LG-Fc, while control peptides lacking the organized charge distribution showed no binding (Fig. 5D-E). To determine whether this domain mediates the interaction between CD300LG and CMs, hCD300LG-Fc was pre-incubated with hA4-Cterm or the hA4- Scr peptide prior to assessing the binding of Dil-CMs. The hA4-Cterm peptide significantly inhibited Dil-CM binding by ∼40-50%, while hA4-Scr had no effect (Fig. 5F). Similarly, hA4-Cterm but not hA4-Scr produced a dose-dependent decrease in Dil-CM binding to TIME cells overexpressing CD300LG (Fig. 5G).

Finally, we assessed the physiological relevance of the ApoA4 c-terminal region in CM metabolism *in vivo*. WT mice treated with the mA4-Cterm peptide exhibited impaired TG clearance during an OFTT compared to mA4-Scr or saline control, recapitulating the phenotype of Cd300lg-KO mice, albeit to a lesser extent (Fig. 5H). Together, these data demonstrate that CD300LG binds directly to CMs via ApoA4 to facilitate their clearance, and that the repeating EQXQ motif within the c-terminus of ApoA4 is at least partially responsible for this binding activity.

## Discussion

Previous studies have reported genetic associations between the R82C variant of CD300LG and altered plasma lipid levels; however, the biological function of CD300LG and its direct involvement in lipid metabolism remained largely undefined.^9–15^ In this study, we identified CD300LG as a novel receptor of TRLs that binds directly to ApoA4 and facilitates TRL clearance at the microvascular endothelium. Our findings demonstrate that CD300LG deficiency in mice significantly impairs postprandial lipid metabolism and results in elevated plasma TG levels specifically under conditions of dietary lipid excess. This discovery elucidates a previously unknown mechanism underlying the genetic association between CD300LG variants and dyslipidemia. Furthermore, our human genetic data pinpointed the effect of CD300LG on TRLs and demonstrated a causal relationship whereby reductions in CD300LG protein levels lead to increased TG levels and CAD risk in humans. Collectively, these data establish CD300LG as a key receptor for TRLs, facilitating their processing at the vascular endothelium.

The CD300 family has been largely studied for its role in the immune system, and CD300LG has been described as a ligand for L-selectin.^5^ Most of the CD300 family members are highly expressed within immune cells with little to no expression in other cell types.^6^ CD300LG, on the other hand, is expressed in microvascular ECs rather than immune cells, and its gene locus is also distinct from the remaining CD300 family members.^6,7^ CD300LG is further distinguished by its mucin domain and its lack of functional intracellular signaling domains seen on other CD300 family members.^6^ Altogether, this supports the idea that CD300LG may play a very different role from its paralogs.

Chylomicrons produced by the small intestine following a meal are processed by LPL to enable the uptake of lipids primarily within muscle, heart and adipose. The loss of LPL, or proteins which directly regulate its localization or activity, all result in significant changes to plasma TGs in both the fasted and fed state.^26,27,30,33,34^ In contrast, Cd300lg-KO mice maintained normal TG levels under fasting conditions or on chow diet, yet showed a profound defect in TG clearance after exposure to a large oral lipid load (lipid gavage or high-fat diet feeding). Further, CD300LG does not appear to be essential to the tissue distribution of fat, as plasma TG levels of Cd300lg-KO mice not only returned to normal (albeit delayed) following lipid intake, but there were also no differences in tissue weights for adipose, liver or skeletal muscle. This differs from mice lacking direct regulators of LPL such as GPIHBP1 or ANGPTL8, which exhibit significant alterations in both weight gain and adiposity.^42,43^ This distinct phenotype is consistent with our findings that the loss of CD300LG did not directly affect vascular luminal LPL activity or protein levels. Of note, *Cd300lg* does not appear to be transcriptionally regulated by feeding (Fig. S2B) or lipid gavage (Fig. S6E). Therefore, we believe that CD300LG provides a unique additional regulatory mechanism for plasma TRL metabolism, which is dispensable for baseline plasma TRL catabolism in mice but becomes crucial when the body is confronted with an acute influx of dietary lipid.

TRL margination along the vascular endothelium has been attributed to its interaction with the LPL- GPIHBP1 complex and heparan sulfate proteoglycans.^35,36^ Our findings introduce CD300LG as an additional contributor to this process. Biochemical studies suggested recombinant CD300LG was capable of binding TRLs directly. In addition, overexpression of CD300LG alone was sufficient for CM and VLDL binding to TIME cells, which do not express GPIHBP1 or LPL. While the ability of the GPIHBP1-LPL complex to provide high-affinity binding to TRLs likely explains why Cd300lg-KO mice exhibit normal plasma TG levels under low lipid conditions, our data suggest that CD300LG serves as an additional binding mechanism for TRLs to the endothelium via ApoA4 to enhance TRL clearance under high lipid conditions (Fig. 6). We therefore propose that CD300LG binding to ApoA4-containing TRLs may serve as a low affinity, but high-capacity binding system to enhance the LPL processing specifically of postprandial lipids.

**Figure 6.**
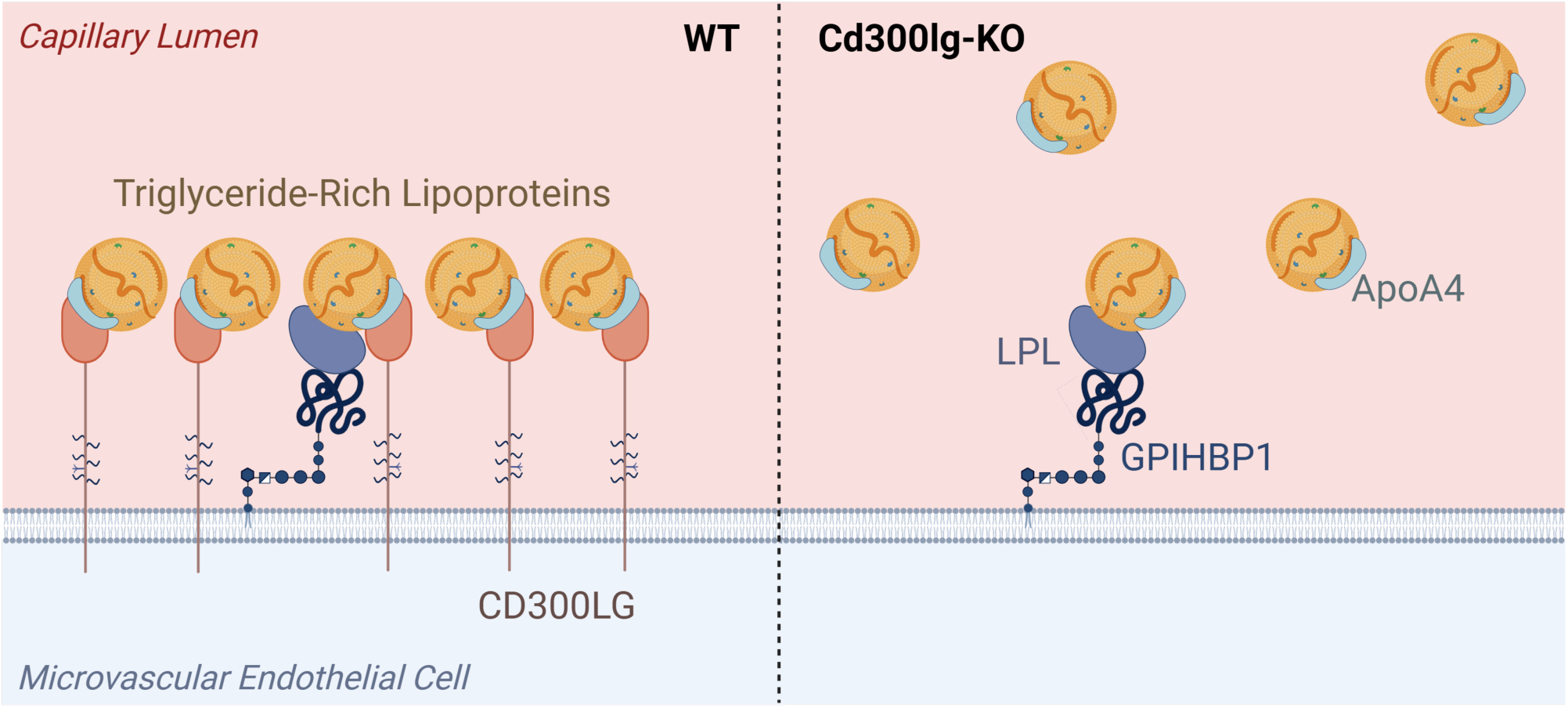
Proposed model for CD300LG in TRL metabolism. The GPIHBP1-LPL complex enables high affinity binding of TRLs that is sufficient to enable their processing by LPL under low-lipid conditions. Under high lipid postprandial conditions, CD300LG provides additional binding sites for TRLs on the microvascular endothelium to enhance processing by LPL. The loss of CD300LG under high-lipid conditions results in insufficient binding sites for TRLs on ECs, a reduced rate of TRL metabolism, and hypertriglyceridemia.

While ApoA4 is found on both VLDL and CMs, it is primarily produced by the small intestine and is involved in both the assembly and secretion of nascent CMs.^38,39^ Previous studies have indicated that ApoA4 has a multifaceted role in both CM biogenesis and metabolism. The plasma TG phenotype from the available ApoA4 knockout mouse line is somewhat complicated by the fact that there is a significant reduction in the expression of the LPL regulator ApoC3 in the ApoA4 knockout mice.^44^ That being said, CMs isolated from ApoA4 knockout mice display significantly impaired clearance when injected into wild type mice, consistent with our findings that ApoA4 serves as a ligand for CD300LG to enable TRL binding and processing.^45^

Structural analysis of ApoA4 suggested the C-terminus of ApoA4 as a potential protein-protein binding site.^41^ To this end, the 16-amino acid repeating motif (aa 374 – 389) of ApoA4 in isolation demonstrated saturable binding to CD300LG. Further, this peptide could also compete with CMs for binding to CD300LG, suggesting that this interaction is critical for CD300LG-mediated TRL binding. It is worth noting that ApoA4 is an exchangeable apolipoprotein, and significant amounts of ApoA4 are found both as part of the HDL fraction and as free ApoA4 within plasma. It is not clear why free ApoA4 or ApoA4 found in HDL do not compete for binding to CD300LG, however biochemical studies of lipid-free ApoA4 have provided a possible explanation. While no crystal structure is available for the C-terminus of ApoA4, cross-linking studies have shown that the N- and C-termini of ApoA4 closely interact in the absence of lipid, regardless of whether ApoA4 is in its monomeric or dimeric form.^46^ We hypothesize that ApoA4 incorporated into TRLs may present an exposed C-terminus resulting in a higher affinity to CD300LG compared to its lipid-free form. Additional studies are needed to determine exactly how ApoA4 in these different fractions interacts with CD300LG.

In conclusion, our studies identified CD300LG as an endothelial TRL receptor that plays a crucial role in postprandial TRL clearance. This discovery reveals a previously unrecognized regulatory mechanism in plasma lipid homeostasis. Elevated postprandial TRLs are increasingly recognized as major contributors to the residual risk of CAD that persists even after LDL-cholesterol is well controlled, yet there are limited therapeutics specifically targeting this risk factor.^2,3^ Our discovery of CD300LG’s role in postprandial TRL metabolism fills a critical gap in understanding how the body handles postprandial lipemia. By promoting the efficient clearance of postprandial TRLs, CD300LG may help mitigate the residual CAD risk caused by prolonged TRL circulation. Taken together, these data provide a deeper understanding of postprandial lipid homeostasis and open exciting new avenues for pharmacological intervention of postprandial lipid metabolism for the treatment of cardiometabolic diseases.

### Limitations of the study

While our study identifies a key function for CD300LG in TRL metabolism, several limitations should be acknowledged. First, the cis-pQTLs identified in this study were determined based on plasma protein levels of CD300LG with the assumption that the circulating CD300LG level correlates with the cell surface CD300LG level. The exact mechanism of how CD300LG is released into circulation and its potential biological function needs further study. Further, it should be noted that the lipoprotein profiles measured by NMR in UK Biobank are separated into fractions based on particle diameter, and thus these data cannot differentiate between VLDL and CMs.^21,22^ Next, while we identified the ApoA4 c-terminal region is essential for CD300LG binding, other domains in ApoA4 or other additional apolipoproteins may also contribute to the binding and remain to be investigated. Finally, whether CD300LG influences TRL metabolism through additional mechanisms beyond direct binding, such as facilitating apolipoprotein transfer, was not explored in depth. Future studies examining these possibilities will help clarify the full scope of CD300LG’s involvement in lipid metabolism.

## Resource Availability

### Lead Contact

For further information and questions or requests regarding resources, contact Xin Rong.

### Materials Availability

All unique reagents and materials used in this study are available from the vendors listed. Complete details for ordering the reagents will be provided upon request. The availability of the Cd300lg-KO mouse line generated in this study is restricted and will require a materials transfer agreement prior to distribution.

### Data and Code Availability

All large data sets used in this study were obtained from publicly available resources and are acknowledged in the methods. No original code is reported by this paper. Additional information required to reanalyze the data reported in this paper is available from the lead contact upon request.

## Acknowledgments

We would like to acknowledge the efforts of Mackenzie Marshall, Cyndi Pinkus, Shannon Kalsow, Elizabeth Thomason, Rowann Mostafa, Jake Delmore, Eliza Bollinger, Amanda Audesse, Kerry Kelleher and Ruth Elgamal for technical assistance, and William Sessa for discussions and advice on the manuscript.

The human genetic analyses using UK Biobank Pharma Proteomics Project data were conducted under application 65851. The CAD GWAS summary statistics from Million Veteran Program used in the current publication are based on the study data downloaded from the dbGaP website, under phs001672.v8.p1. We acknowledge the use of GWAS summary statistics for lipoprotein profiles from MRC IEU OpenGWAS database.^47^ We thank the participants and investigators of UK Biobank, FinnGen, Million Veteran Program, Global Lipids Genetics Consortium, and CardiogramC4Dplus study.

CytoReason performed the reprocessing and annotation of the human skeletal muscle scRNAseq dataset.

Graphics were created with https://BioRender.com. Helical wheel diagrams were generated using the Net Wheels tool (https://neutrophil-proteome.shinyapps.io/netwheels/).^48^ Bulk tissue gene expression data (median gene-level TPM by tissue) was obtained from the GTEx Portal on 05/06/2025.

## Author contributions

M.E.G, K.M.T, R.J.R.F, and X.R. conceived the project and designed the experiments. M.E.G and K.M.T. performed the experiments and analyzed the data. H.K. performed genetic analyses. A.R.Y, K.T.N., N.A., M.M., A.W., D.A., R.S., and B.L. contributed to *in vivo* experiments. J.F. performed flow cytometry analyses.

Y.A. led the design and generation of Cd300lg-KO mice. J.C. performed proteomics analysis of TRLs. M.P. re-analyzed single cell RNAseq data. M.E.G., K.M.T., H.K., R.J.R.F., and X.R. wrote the manuscript with input from all authors.

## Declaration of interests

All authors of this manuscript are current employees of Pfizer Inc., or were employees of Pfizer Inc. at the time when the studies were conducted.

## Supplemental information

Supplemental Figures. Figures S1 – S6

Supplemental Tables. Tables S1 – S4

## Methods

### Materials

TIME cells were obtained from ATCC (CRL4025) and cultured in vascular cell basal medium (ATCC, PCS- 100-030) supplemented with microvascular endothelial cell growth kit-VEGF (ATCC, PCS-110-041) with additional FBS to a final concentration of 10%. qPCR primers and TaqMan probes were obtained from ThermoFisher. Peptides matching the mouse or human c-terminus sequences of ApoA4 or scrambled control peptides were custom ordered with an N-terminal biotin tag from CPC Scientific. Empty vector or human CD300LG-expressing adenoviruses were custom ordered from Vector Biolabs in the Ad-CMV-intron- myc-DDK backbone.

### Mice

All animal experiments were conducted following study protocols and procedures reviewed and approved by Pfizer Institutional Animal Care and Use Committee. The facilities that supported this work are fully accredited by AAALAC International.

Mice were either group-housed or housed individually (metabolic phenotyping studies, Fig. 3, S3, S4) in Innovive cages (Innorack IVC Mouse 3.5) under a standard 12-h light/12-h dark cycle (06:00 h: 18:00 h) in a temperature- and humidity-controlled environment at room temperature (22 ± 1℃). Mice were given *ad libitum* access to tap water and standard chow (Inotiv Teklad 2916; Inotiv, Inc., West Lafayette, IN) or 60% high-fat diet (Research Diets D12492, New Brunswick, NJ). For metabolic studies, 6-week-old mice were placed on chow or high-fat diet for 18 weeks and assessed at the indicated time points. Body composition was assessed using the EchoMRI 4-in-1 500 Body Composition Analyzer (EchoMRI, Houston, TX). Tissues were collected from either overnight fasted or *ad libitum* fed mice after 18-weeks of diet and tissue weights were measured prior to snap freezing for further analysis. For fasted and refed analyses, mice were fasted overnight with *ad libitum* access to water and refed with either chow or high-fat diet for 2 hours.

### Generation of Cd300lg knockout mice

Mice deficient for the Cd300lg gene were created and maintained on the C57BL/6J strain (JAX, #000664) utilizing CRISPR/Cas9 technology.^49^ Knockout of the gene was achieved by introducing frame- shifting small insertions or deletions (indels) into exon 2. Cas9 guides were designed for efficient editing while minimizing potential risk of off-target editing (www.benchling.com). The seed sequence of the Cas9 sgRNA used for editing was: 5’AGGAAGTATTGGTGCCGGCA. To generate F0 mice, 1-cell embryos were harvested from superovulated females and electroporated with an *in vitro* synthesized sgRNA complexed with Cas9 protein (IDT, #1081058). Manipulated embryos were then implanted into pseudo-pregnant females. F0 mice were genotyped by PCR amplification with primers, 5’GCCAGCCTGACTTGTGTCTC and 5’ATGGCTCAGATGCTCTCACC, followed by Sanger sequencing. Raw sequencing data were analyzed with the TIDE software (https://tide.nki.nl/) to estimate frequency of individual indels in each animal. Sequences revealed that selected F0 mice edited with sgRNA #485 contained a 4-bp deletion within exon 2 that resulted in a premature stop codon (Fig. S2A). To establish a colony, selected F0 mice were crossed with wild-type C57BL/6J mice for germline transmission and heterozygous progeny were further bred to produce littermate-matched wildtype and homozygous knockout cohorts.

### Human genetic analyses

GWAS summary statistics for NMR-based plasma lipoprotein and lipid measurements in UK Biobank were obtained from IEU OpenGWAS project (https://gwas.mrcieu.ac.uk/).^47^ GWAS summary statistics for blood lipid levels were obtained from the Global Lipids Genetics Consortium (GLGC).^18^ GWAS summary statistics for plasma CD300LG protein measurement were obtained from the UK Biobank Pharma Proteomics Project (UKB-PPP).^20^ Meta-analysis for CAD was performed under the fixed-effect model using METAL across GWAS summary statistics from CARDIoGRAMplusC4D^23^, Million Veteran Program (MVP)^24^, and FinnGen (release 12)^25^ studies. Fine-mapping of cis-pQTL signals and colocalization were performed using the runsusie and coloc functions of coloc package (v.5.1.0). The imputed genotype data from the European participants of UK Biobank were used as the LD reference. For Mendelian randomization analyses, 8 lead variants of the independent cis-pQTLs were used as the instrumental variables (IVs). When the IV variants were not available in the summary statistics of the outcome trait, variants in LD (r^2^>0.8) were used instead. GCTA-COJO (v.1.25.2) was used to derive the joint effects among the IVs for both the exposure and the outcome traits. Mendelian randomization analysis was performed using TwoSampleMR (v.0.5.10) and MendelianRadomizaton (v.0.10.0) packages.

### Quantitative RT-PCR

Animals were euthanized via CO2 and tissues were immediately dissected and snap frozen in liquid nitrogen. Ribonucleic acid (RNA) was extracted with TRIzol reagent (Invitrogen, #15596026) and purified using RNeasy RNA Isolation Kit (QIAGEN, Germantown, MD) then reverse transcribed into complementary deoxyribonucleic acid (cDNA) with a High-Capacity cDNA Reverse Transcription Kit (Applied Biosystems, Foster City, CA). Quantitative reverse transcriptase polymerase chain reaction (qRT-PCR) was performed using TaqMan reagents and primer-probes (Applied Biosystems, Foster City, CA). For liver, eWAT, gastrocnemius, and heart, gene expression was normalized to the average of the control genes cyclophilin A (Ppia) and 14-3-3zeta (Ywhaz). For small intestine, gene expression was normalized to the average of the control genes TATA box binding protein (Tbp) and ribosomal protein lateral stalk subunit P0 (Rplp0).

### Flow Cytometry

To assess the expression of CD300LG protein in endothelial cells, WT and Cd300lg-KO mouse hearts were isolated and digested in Hank’s Balanced Salt Solution (Gibco, 14025-092) containing 2.5% BSA (Sigma-Aldrich, A7030), 4 mg/mL Collagenase A (Roche, 11088793001), 0.2 mg/mL Collagenase XI (Sigma-Aldrich, C7652), 0.12 mg/mL DNAse I (Sigma-Aldrich, DN25), and 0.4 mg/mL Hyaluronidase (Sigma-Aldrich, HX0514- 1) for 45 minutes at 37℃. The single cell suspension was filtered through a 40 µm strainer and suspended in RBC lysis buffer for 5 minutes. The cells were then suspended in FACS buffer (PBS containing 2% FBS), blocked with anti-mouse CD16/32 antibody (Biolegend, 101301) and stained with a 1:100 dilution of antibodies against CD31 (Biolegend, 102410), CD45 (Biolegend, 103108), and either CD54 (Biolegend, 116107) or CD300LG (Biolegend, 147104) along with a viability dye (ThermoFisher, 65-0863-18).

Flow cytometry on samples was carried out on a BD LSR Fortessa A in the Pfizer Flow Cytometry Technology Center. Compensation was carried out using single stains conjugated to UltraComp eBeads (Invitrogen, 01222242) and a single stain of viability dye using heart cells in which 40% of the sample was boiled for 3 minutes. Gating was accomplished through analysis of fluorescence minus one (FMO) controls for CD31, CD45, CD54, and CD300LG. Then, data was collected for each sample stopping at either 100,000 CD31+; CD45- events (x2 hearts) or 50,000 CD31+; CD45- events (x2 hearts).

### Assessment of plasma TGs and cholesterol

Plasma TGs were assessed using the Infinity Triglycerides reagent (Thermo Fisher, #TR22421) according to the manufacturer’s protocol. TG values were calculated by comparing to a standard curve prepared with the multi calibrator lipids reagent (Fujifilm, #464-01601). Plasma cholesterol was assessed with the Cholesterol E kit according to the manufacturer’s protocol (Fujifilm, #999-02601).

### Glucose and insulin tolerance tests

Prior to performing glucose and insulin tolerance tests, mice were fasted for 5h. For glucose tolerance, mice were administered a 2 g/kg bodyweight dose of dextrose by oral gavage. For insulin tolerance, mice were administered 0.75 U/kg bodyweight of Humulin by intraperitoneal injection. Blood glucose was monitored from the tail vein at the timepoints shown using an AlphaTrak2 glucometer (Zoetis).

### Oral fat tolerance tests

Mice were fasted overnight before being administered a 6 µL/g bodyweight dose of olive oil (Sigma Aldrich, #O1514) by oral gavage. Tail vein blood was collected for each time point and plasma prepared for the assessment of plasma TG levels. To assess the effects of the mouse ApoA4 c-terminal (mA4-Cterm) and scrambled (mA4-Scr) peptides on TGs during an OFTT, peptides were suspended in sterile physiologic saline at a concentration of 400 µg/mL and baseline bleeds were performed. Mice were then anesthetized with isoflurane and administered with 40 µg peptide/mouse by retroorbital injection. Approximately 15 minutes later, mice were administered 10 µL/g bodyweight of olive oil by oral gavage and TGs were monitored at the indicated time points.

### scRNAseq expression dot plots

The human skeletal muscle single-cell dataset was generated by Turiel et. al. (GEO Accession: GSE235143).^50^ The dataset was reprocessed using Cellbender for removal of ambient RNA, followed by doublet removal using DoubletFinder. Subsequent analysis was conducted using the Seurat workflow.^51–53^ The endothelial cells were subclustered to identify endothelial subtypes.

The human adipose data was generated by Emont et. al. (GEO Accession: GSE176171).^54^ To identify the endothelial subtypes, the clustering resolution of the dataset was adjusted to 1.2 within Seurat, based on the splitting of the endothelial cells into subtypes. Marker genes for the endothelial subtypes (All ECs: VWF, CDH5, Artery ECs: SOX5, Venule ECs: ACKR1, Lymphatic ECs: PROX1, Capillary ECs: RGCC) were used to annotate clusters. Gene expression levels were visualized using Seurat’s DotPlot function.^53^

### VLDL secretion and intestinal absorption assays

Mice were fasted overnight before being administered Poloxamer-407 (P-407) (Sigma Aldrich, #16758) at a dose of 500 mg/kg bodyweight by intraperitoneal injection. For VLDL secretion, plasma TGs were monitored at the indicated time points following P-407 dosing. For intestinal absorption, mice were administered olive oil by oral gavage at a dose of 10 µL/g bodyweight 30 minutes after receiving P-407. Plasma TGs were monitored at the indicated time points following olive oil administration.

### Quantitation of post-heparin plasma LPL mass and activity

LPL enzymatic activity and protein levels were measured from WT and Cd300lg-KO mice maintained on either chow or on HFD for 18 weeks. For chow-fed mice, measurements were made from mice that were either fasted overnight or administered an oral lipid gavage of olive oil (10 µL/g bodyweight) 2 hours prior to the beginning of the experiment. For HFD-fed mice, measurements were made from mice that were either fasted overnight or refed for 2 hours on HFD prior to the experiment. Baseline pre-heparin plasma was collected for all mice prior to the administration of heparin (Sigma Aldrich, #H3393) by retroorbital injection (2 U/g bodyweight in sterile saline). 15 minutes after heparin administration, mice were euthanized by CO_2_ and post-heparin plasma was collected by cardiac puncture. LPL mass was determined with a mouse LPL ELISA kit (Biotang, m7614). Lipase activity was measured in both pre-heparin and post-heparin samples using a fluorimetric kit (Abcam, ab204721) according to the manufacturer’s protocol. LPL activity was calculated by subtracting the activity of the pre-heparin plasma from the activity of the post-heparin plasma.

### Proteomics analysis of triglyceride-rich lipoprotein particles

Plasma was collected from WT and Cd300lg-KO mice at the end of the intestinal absorption assays (4 hours after olive oil gavage in the presence of P-407) and TRLs were isolated by ultracentrifugation by overlaying plasma with an NaCl solution with a density of 1.006 g/mL, and centrifuging in a TLA 110 rotor at 110,000 rpm for 6 hours at 4℃. The top layer of TRLs was collected and dialyzed against PBS with 5 µM EDTA. Isolated chylomicrons were treated with 4% sodium dodecyl sulfate supplemented with 0.1M Tris, pH 8.5. Protein was precipitated overnight at -20°C in the presence of excess acetone. The following day, protein was pelleted by centrifugation and rinsed with fresh acetone. Protein was digested overnight in the presence of trypsin, followed by desalting by solid-phase extraction (C18 Sep-Pak, Waters). Peptide digest was quantified by fluorometric assay (Pierce). For each sample, an aliquot of 1.5μg of peptide material was desalted using a C18 STAGE tip extraction (Empore) prior to analysis using a Vanquish Neo UHPLC system coupled to a Thermo Exploris 480 mass spectrometer. Samples were analyzed in data-independent acquisition mode and searched using Spectronaut 19 (Biognosys). Statistical analysis was performed using in-house scripts and p-values were adjusted by False Discovery Rate.

### Binding of lipoprotein particles to TIME cells

To prepare Dil-labeled CMs, 500 µL of human CMs (Athens Research, 12-16-030825) were mixed with 20 µL of a 3 mg/mL (in DMSO) stock solution of Dil stain (Invitrogen, D3911) and 4 mL of lipoprotein-depleted fetal bovine serum (Kalen Biomedical, 880100-1) and incubated at 37℃ overnight. Labeled CMs were transferred into ultracentrifuge tubes, overlayed with an NaCl solution with a density of 1.006 g/mL, and centrifuged in a TLA 110 rotor at 110,000 rpm for 6 hours at 4℃. The top layer of labeled CMs was collected and dialyzed against PBS with 5 µM EDTA. Final protein concentration of labeled CMs was determined by a BCA assay.

TIME cells were cultured and seeded into 96-well PhenoPlates (Revvity, 6055300) and allowed to attach for 4-6 hours. TIME cells were then infected with either empty vector or human CD300LG-expressing adenovirus and cultured overnight. Lipoprotein particle binding on TIME cells was then carried out as has been previously published with CHO cells by Beigneux *et. al*. with minor modifications.^30^ In brief, TIME cells were washed twice with ice cold CM binding buffer (PBS, 1 mM CaCl_2_, 1 mM MgCl_2_, 0.5% BSA) and incubated with 1 µg/mL Dil-labeled lipoprotein particles as indicated (or other concentrations as indicated) for 2 hours at 4℃. The cells were then washed five times with CM binding buffer and fixed with 4% PFA. The binding of Dil-labeled lipoprotein particles was assessed by high-content fluorescence microscopy using an Opera Phenix Plus and quantified as mean spot area per well normalized by cell number. For competition experiments, pre-incubations were performed as indicated in figure legends.

### Direct binding of chylomicrons to purified CD300LG-Fc

Human CD300LG-Fc (R&D Systems, 10615-NE) or normal human IgG1 isotype control (internal) were attached to a 96-well polystyrene plate (Corning, 3631) by adding 1 µg total protein in 100 µL of PBS per well and incubating overnight at 4℃. The plate was washed 3 times with PBS and blocked with the addition of 200 µL CM binding buffer and incubation at 37℃ for 2 hours. After blocking, the plate was aspirated and 1 µg of Dil-CMs was added to each well suspended in 100 µL of CM binding buffer and incubated for 2 hours at room temperature protected from light. The plate was washed 5 times with PBS, and each well was imaged in its entirety using an Opera Phenix Plus. The binding of Dil-CMs was quantified by calculating the sum total intensity within the Dil channel for each well.

### Direct binding of purified CD300LG-Fc to apolipoproteins

A total of 25 pmol of each of the recombinant human apolipoproteins ApoC2 (LSBio, LS-G20488), ApoC3 (LSBio, LS-G20489), ApoA4 (Sino Biological, 16082-H08H), and ApoA5 (Prospec-Tany, Cyt-025) were attached to a 96-well polystyrene plate similar to as described for CD300LG-Fc. The plate was washed 3 times with PBS and blocked with the addition of 200 µL CM binding buffer and incubation at 37℃ for 2 hours. After blocking, the plate was aspirated and 25 pmol of CD300LG-Fc or normal human IgG1 isotype control (control Fc, internal) was added to each well in 100 µL of CM binding buffer and incubated at room temperature for 2 hours with shaking (500 rpm). The plate was washed 5 times with PBS and 100 µL of goat anti-human Ig-HRP detection antibody (Southern Biotech, 2081-05) was added to each well at a 1:10,000 dilution in CM binding buffer. The plate was incubated with detection antibodies for 1 hour at room temperature with shaking. The plate was washed 5 times with PBS and 100 µL of TMB substrate (ThermoFisher, 34022) was added and incubated for 10 minutes at room temperature before the addition of 50 µL of stop solution (ThermoFisher, N600). Absorbance was measured at 450 nm and corrected for absorbance at 540 nm. The percentage of total protein bound was calculated by comparing to a standard curve prepared by binding either control Fc or CD300LG-Fc directly to the plate. For the competition experiments with hA4-Cterm or hA4-Scr, CD300LG-Fc coated wells were pre-incubated with 10 µM hA4- Cterm or hA4-Scr, or CM binding buffer alone for 2 hours at room temperature before the addition of Dil- CMs.

### Direct binding of ApoA4 c-terminal peptides to purified CD300LG-Fc

Human C300LG-Fc was attached to a 96-well polystyrene plate at 10 pmol per well as described above. The wells were then incubated with the peptides at the concentrations indicated (100 µL per well) for 2 hours at room temperature with shaking. The plate was washed 5 times with CM binding buffer and the peptides were detected via the N-terminal biotin tag using Streptavidin-HRP (Invitrogen, 434323) at a 1:2500 dilution and incubated at 37℃ for 10 minutes. The plate was washed 5 times with CM binding buffer and 100 µL of TMB substrate was added and incubated for 15 minutes at 37℃ followed by the addition of 50 µL of stop solution. Absorbance was read at 450 nm and corrected for absorbance at 540 nm.

### Quantification and statistical analysis

Sample size is indicated by individually plotted points or indicated within the figure legend where individual points are not plotted. Error bars denote SEM for *in vivo* experiments and SD for *in vitro* experiments. Significance was calculated by t-test, one-way or two-way ANOVA as appropriate using GraphPad Prism software.

## Notes

### Funding

- Pfizer (United States)

## References

1. Gugliucci, A. (2023). The chylomicron saga: time to focus on postprandial metabolism. Front Endocrinol (Lausanne) 14, 1322869. 10.3389/fendo.2023.1322869.

2. Abol-eineen, M.W., Mostafa, T.M., Salama, A.E., and Zigheel, H. A. k. F. (2021). Significance of postprandial triglycerides in coronary artery disease. Zagazig University Medical Journal 0, 20–28. 10.21608/zumj.2019.11200.1176.

3. Patsch, J.R., Miesenböck, G., Hopferwieser, T., Mühlberger, V., Knapp, E., Dunn, J.K., Gotto Jr., A.M., and Patcsh, W. (1992). Relation of Triglyceride Metabolism and Coronary Artery Disease. Studies in the Postprandial State. Arteriosclerosis and Thrombosis 12, 1336–1345.

4. Umemoto, E., Takeda, A., Jin, S., Luo, Z., Nakahogi, N., Hayasaka, H., Lee, C.M., Tanaka, T., and Miyasaka, M. (2013). Dynamic changes in endothelial cell adhesion molecule nepmucin/CD300LG expression under physiological and pathological conditions. PLoS One 8, e83681. 10.1371/journal.pone.0083681.

5. Umemoto, E., Tanaka, T., Kanda, H., Jin, S., Tohya, K., Otani, K., Matsutani, T., Matsumoto, M., Ebisuno, Y., Jang, M.H., et al. (2006). Nepmucin, a novel HEV sialomucin, mediates L-selectin-dependent lymphocyte rolling and promotes lymphocyte adhesion under flow. J Exp Med 203, 1603–1614. 10.1084/jem.20052543.

6. Borrego, F. (2013). The CD300 molecules: an emerging family of regulators of the immune system. Blood 121, 1951–1960. 10.1182/blood-2012-09-435057.

7. Takatsu, H., Hase, K., Ohmae, M., Ohshima, S., Hashimoto, K., Taniura, N., Yamamoto, A., and Ohno, H. (2006). CD300 antigen like family member G: A novel Ig receptor like protein exclusively expressed on capillary endothelium. Biochem Biophys Res Commun 348, 183–191. 10.1016/j.bbrc.2006.07.047.

8. Jin, S., Umemoto, E., Tanaka, T., Shimomura, Y., Tohya, K., Kunizawa, K., Yang, B.G., Jang, M.H., Hirata, T., and Miyasaka, M. (2008). Nepmucin/CLM-9, an Ig domain-containing sialomucin in vascular endothelial cells, promotes lymphocyte transendothelial migration in vitro. FEBS Lett 582, 3018–3024. 10.1016/j.febslet.2008.07.041.

9. Albrechtsen, A., Grarup, N., Li, Y., Sparso, T., Tian, G., Cao, H., Jiang, T., Kim, S.Y., Korneliussen, T., Li, Q., et al. (2013). Exome sequencing-driven discovery of coding polymorphisms associated with common metabolic phenotypes. Diabetologia 56, 298–310. 10.1007/s00125-012-2756-1.

10. Surakka, I., Horikoshi, M., Magi, R., Sarin, A.P., Mahajan, A., Lagou, V., Marullo, L., Ferreira, T., Miraglio, B., Timonen, S., et al. (2015). The impact of low-frequency and rare variants on lipid levels. Nat Genet 47, 589–597. 10.1038/ng.3300.

11. Helgadottir, A., Gretarsdottir, S., Thorleifsson, G., Hjartarson, E., Sigurdsson, A., Magnusdottir, A., Jonasdottir, A., Kristjansson, H., Sulem, P., Oddsson, A., et al. (2016). Variants with large effects on blood lipids and the role of cholesterol and triglycerides in coronary disease. Nat Genet 48, 634–639. 10.1038/ng.3561.

12. Metz, S., Krarup, N.T., Bryrup, T., Stoy, J., Andersson, E.A., Christoffersen, C., Neville, M.J., Christiansen, M.R., Jonsson, A.E., Witte, D.R., et al. (2022). The Arg82Cys Polymorphism of the Protein Nepmucin Implies a Role in HDL Metabolism. J Endocr Soc 6, bvac034. 10.1210/jendso/bvac034.

13. Wang, Z., Chen, H., Bartz, T.M., Bielak, L.F., Chasman, D.I., Feitosa, M.F., Franceschini, N., Guo, X., Lim, E., Noordam, R., et al. (2020). Role of Rare and Low-Frequency Variants in Gene-Alcohol Interactions on Plasma Lipid Levels. Circ Genom Precis Med 13, e002772. 10.1161/CIRCGEN.119.002772.

14. Liu, D.J., Peloso, G.M., Zhan, X., Holmen, O.L., Zawistowski, M., Feng, S., Nikpay, M., Auer, P.L., Goel, A., Zhang, H., et al. (2014). Meta-analysis of gene-level tests for rare variant association. Nat Genet 46, 200–204. 10.1038/ng.2852.

15. Kanoni, S., Masca, N.G., Stirrups, K.E., Varga, T.V., Warren, H.R., Scott, R.A., Southam, L., Zhang, W., Yaghootkar, H., Muller-Nurasyid, M., et al. (2016). Analysis with the exome array identifies multiple new independent variants in lipid loci. Hum Mol Genet 25, 4094–4106. 10.1093/hmg/ddw227.

16. Karjalainen, M.K., Karthikeyan, S., Oliver-Williams, C., Sliz, E., Allara, E., Fung, W.T., Surendran, P., Zhang, W., Jousilahti, P., Kristiansson, K., et al. (2024). Genome-wide characterization of circulating metabolic biomarkers. Nature 628, 130–138. 10.1038/s41586-024-07148-y.

17. Cannon, J.P., O’Driscoll, M., and Litman, G.W. (2012). Specific lipid recognition is a general feature of CD300 and TREM molecules. Immunogenetics 64, 39–47. 10.1007/s00251-011-0562-4.

18. Graham, S.E., Clarke, S.L., Wu, K.H., Kanoni, S., Zajac, G.J.M., Ramdas, S., Surakka, I., Ntalla, I., Vedantam, S., Winkler, T.W., et al. (2021). The power of genetic diversity in genome-wide association studies of lipids. Nature 600, 675–679. 10.1038/s41586-021-04064-3.

19. Klarin, D., Damrauer, S.M., Cho, K., Sun, Y.V., Teslovich, T.M., Honerlaw, J., Gagnon, D.R., DuVall, S.L., Li, J., Peloso, G.M., et al. (2018). Genetics of blood lipids among ∼300,000 multi-ethnic participants of the Million Veteran Program. Nat Genet 50, 1514–1523. 10.1038/s41588-018-0222-9.

20. Sun, B.B., Chiou, J., Traylor, M., Benner, C., Hsu, Y.H., Richardson, T.G., Surendran, P., Mahajan, A., Robins, C., Vasquez-Grinnell, S.G., et al. (2023). Plasma proteomic associations with genetics and health in the UK Biobank. Nature 622, 329–338. 10.1038/s41586-023-06592-6.

21. Julkunen, H., Cichonska, A., Tiainen, M., Koskela, H., Nybo, K., Makela, V., Nokso-Koivisto, J., Kristiansson, K., Perola, M., Salomaa, V., et al. (2023). Atlas of plasma NMR biomarkers for health and disease in 118,461 individuals from the UK Biobank. Nat Commun 14, 604. 10.1038/s41467-023-36231-7.

22. Borges, M.C., Haycock, P.C., Zheng, J., Hemani, G., Holmes, M.V., Davey Smith, G., Hingorani, A.D., and Lawlor, D.A. (2022). Role of circulating polyunsaturated fatty acids on cardiovascular diseases risk: analysis using Mendelian randomization and fatty acid genetic association data from over 114,000 UK Biobank participants. BMC Med 20, 210. 10.1186/s12916-022-02399-w.

23. Aragam, K.G., Jiang, T., Goel, A., Kanoni, S., Wolford, B.N., Atri, D.S., Weeks, E.M., Wang, M., Hindy, G., Zhou, W., et al. (2022). Discovery and systematic characterization of risk variants and genes for coronary artery disease in over a million participants. Nat Genet 54, 1803–1815. 10.1038/s41588-022-01233-6.

24. Tcheandjieu, C., Zhu, X., Hilliard, A.T., Clarke, S.L., Napolioni, V., Ma, S., Lee, K.M., Fang, H., Chen, F., Lu, Y., et al. (2022). Large-scale genome-wide association study of coronary artery disease in genetically diverse populations. Nat Med 28, 1679–1692. 10.1038/s41591-022-01891-3.

25. Kurki, M.I., Karjalainen, J., Palta, P., Sipila, T.P., Kristiansson, K., Donner, K.M., Reeve, M.P., Laivuori, H., Aavikko, M., Kaunisto, M.A., et al. (2023). FinnGen provides genetic insights from a well-phenotyped isolated population. Nature 613, 508–518. 10.1038/s41586-022-05473-8.

26. Weinstock, P.H., Bisgaier, C.L., Aalto-Setala, K., Radner, H., Ramakrishnan, R., Levak-Frank, S., Essenburg, A.D., Zechner, R., and Breslow, J.L. (1995). Severe Hypertriglyceridemia, Reduced High Density Lipoprotein, and Neonatal Death in Lipoprotein Lipase Knockout Mice. J Clin Invest 96, 2555–2568.

27. Coleman, T., Seip, R.L., Gimble, J.M., Lee, D., Maeda, N., and Semenkovich, C.F. (1995). COOH-terminal disruption of lipoprotein lipase in mice is lethal in homozygotes, but heterozygotes have elevated triglycerides and impaired enzyme activity. J Biol Chem 270, 12518–12525. 10.1074/jbc.270.21.12518.

28. Ghosh, S.S., Wang, J., and Ghosh, S. (2022). Measurement of In Vivo VLDL and Chylomicron Secretion. In Non-Alcoholic Steatohepatitis: Methods and Protocols, D. Sarkar, ed. (Springer US), pp. 63–71. 10.1007/978-1-0716-2128-8_6.

29. Eckel, R.H. (1989). Lipoprotein lipase: a multifunctional enzyme relevant to common metabolic diseases. The New England Journal of Medicine 320, 1060–1068. 10.1056/NEJM198904203201607.

30. Beigneux, A.P., Davies, B.S., Gin, P., Weinstein, M.M., Farber, E., Qiao, X., Peale, F., Bunting, S., Walzem, R.L., Wong, J.S., et al. (2007). Glycosylphosphatidylinositol-anchored high-density lipoprotein-binding protein 1 plays a critical role in the lipolytic processing of chylomicrons. Cell Metab 5, 279–291. 10.1016/j.cmet.2007.02.002.

31. Davies, B.S., Beigneux, A.P., Barnes, R.H., 2nd, Tu, Y., Gin, P., Weinstein, M.M., Nobumori, C., Nyren, R., Goldberg, I., Olivecrona, G., et al. (2010). GPIHBP1 is responsible for the entry of lipoprotein lipase into capillaries. Cell Metab 12, 42–52. 10.1016/j.cmet.2010.04.016.

32. Davies, B.S.J., Goulbourne, C.N., Barnes, R.H., Turlo, K.A., Gin, P., Vaughan, S., Vaux, D.J., Bensadoun, A., Beigneux, A.P., Fong, L.G., and Young, S.G. (2012). Assessing mechanisms of GPIHBP1 and lipoprotein lipase movement across endothelial cells. Journal of Lipid Research 53, 2690–2697. 10.1194/jlr.M031559.

33. Gao, M., Yang, C., Wang, X., Guo, M., Yang, L., Gao, S., Zhang, X., Ruan, G., Li, X., Tian, W., et al. (2020). ApoC2 deficiency elicits severe hypertriglyceridemia and spontaneous atherosclerosis: A rodent model rescued from neonatal death. Metabolism 109, 154296. 10.1016/j.metabol.2020.154296.

34. Sakurai, T., Sakurai, A., Vaisman, B.L., Amar, M.J., Liu, C., Gordon, S.M., Drake, S.K., Pryor, M., Sampson, M.L., Yang, L., et al. (2016). Creation of Apolipoprotein C-II (ApoC-II) Mutant Mice and Correction of Their Hypertriglyceridemia with an ApoC-II Mimetic Peptide. J Pharmacol Exp Ther 356, 341–353. 10.1124/jpet.115.229740.

35. Gin, P., Beigneux, A.P., Voss, C., Davies, B.S., Beckstead, J.A., Ryan, R.O., Bensadoun, A., Fong, L.G., and Young, S.G. (2011). Binding preferences for GPIHBP1, a glycosylphosphatidylinositol-anchored protein of capillary endothelial cells. Arterioscler Thromb Vasc Biol 31, 176–182. 10.1161/ATVBAHA.110.214718.

36. Goulbourne, C.N., Gin, P., Tatar, A., Nobumori, C., Hoenger, A., Jiang, H., Grovenor, C.R., Adeyo, O., Esko, J.D., Goldberg, I.J., et al. (2014). The GPIHBP1-LPL complex is responsible for the margination of triglyceride-rich lipoproteins in capillaries. Cell Metab 19, 849–860. 10.1016/j.cmet.2014.01.017.

37. Jin, H., Zhang, C., Zwahlen, M., von Feilitzen, K., Karlsson, M., Shi, M., Yuan, M., Song, X., Li, X., Yang, H., et al. (2023). Systematic transcriptional analysis of human cell lines for gene expression landscape and tumor representation. Nat Commun 14, 5417. 10.1038/s41467-023-41132-w.

38. Green, P.H.R.G., Robert M.; Riley, John W.; Quinet, Elaine (1980). Human Apolipoprotein A-IV. Intestinal origin and distribution in plasma. J Clin Invest 65, 911–919. 10.1172/JCI109745.

39. Kalogeris, T.J., Rodriguez, M.D., and Tso, P. (1997). Control of synthesis and secretion of intestinal apolipoprotein A-IV by lipid. J Nutr 127, 537S–543S. 10.1093/jn/127.3.537S.

40. VerHague, M.A., Cheng, D., Weinberg, R.B., and Shelness, G.S. (2013). Apolipoprotein A-IV expression in mouse liver enhances triglyceride secretion and reduces hepatic lipid content by promoting very low density lipoprotein particle expansion. Arterioscler Thromb Vasc Biol 33, 2501–2508. 10.1161/ATVBAHA.113.301948.

41. Tubb, M.R., Silva, R., Pearson, K.J., Tso, P., Liu, M., and Davidson, W.S. (2007). Modulation of apolipoprotein A-IV lipid binding by an interaction between the N and C termini. J Biol Chem 282, 28385–28394. 10.1074/jbc.M704070200.

42. Weinstein, M.M., Goulbourne, C.N., Davies, B.S., Tu, Y., Barnes, R.H., 2nd, Watkins, S.M., Davis, R., Reue, K., Tontonoz, P., Beigneux, A.P., et al. (2012). Reciprocal metabolic perturbations in the adipose tissue and liver of GPIHBP1-deficient mice. Arterioscler Thromb Vasc Biol 32, 230–235. 10.1161/ATVBAHA.111.241406.

43. Wang, Y., Quagliarini, F., Gusarova, V., Gromada, J., Valenzuela, D.M., Cohen, J.C., and Hobbs, H.H. (2013). Mice lacking ANGPTL8 (Betatrophin) manifest disrupted triglyceride metabolism without impaired glucose homeostasis. Proc Natl Acad Sci U S A 110, 16109–16114. 10.1073/pnas.1315292110.

44. Weinstock, P.H., Bisgaier, C.L., Hayek, T., Aalto-Setala, K., Sehayek, E., Wu, L., Sheiffele, P., Merkel, M., Essenburg, A.D., and Breslow, J.L. (1997). Decreased HDL cholesterol levels but normal lipid absorption, growth, and feeding behavior in apolipoprotein A-IV knockout mice. Journal of Lipid Research 38, 1782–1794. 10.1016/s0022-2275(20)37153-4.

45. Kohan, A.B., Wang, F., Li, X., Bradshaw, S., Yang, Q., Caldwell, J.L., Bullock, T.M., and Tso, P. (2012). Apolipoprotein A-IV regulates chylomicron metabolism-mechanism and function. Am J Physiol Gastrointest Liver Physiol 302, G628–636. 10.1152/ajpgi.00225.2011.

46. Walker, R.G., Deng, X., Melchior, J.T., Morris, J., Tso, P., Jones, M.K., Segrest, J.P., Thompson, T.B., and Davidson, W.S. (2014). The structure of human apolipoprotein A-IV as revealed by stable isotope-assisted cross-linking, molecular dynamics, and small angle x-ray scattering. J Biol Chem 289, 5596–5608. 10.1074/jbc.M113.541037.

47. Elsworth, B., Lyon, M., Alexander, T., Liu, Y., Matthews, P., Hallett, J., Bates, P., Palmer, T., Haberland, V., Smith, G.D., et al. (2020). The MRC IEU OpenGWAS data infrastructure. bioRxiv, 2020.2008.2010.244293. 10.1101/2020.08.10.244293.

48. Mól, A., Castro, M., and Fontes, W. (2024). NetWheels: A Web Application to Create High Quality Peptide Helical Wheel and Net Projections. Journal of Bioinformatics and Systems Biology 7. 10.26502/jbsb.5107082.

49. Wang, H., Yang, H., Shivalila, C.S., Dawlaty, M.M., Cheng, A.W., Zhang, F., and Jaenisch, R. (2013). One-Step Generation of Mice Carrying Mutations in Multiple Genes by CRISPR/Cas-Mediated Genome Engineering. Cell 153, 910–918. 10.1016/j.cell.2013.04.025.

50. Turiel, G., Desgeorges, T., Masschelein, E., Birrer, M., Zhang, J., Engelberger, S., and De Bock, K. (2023). 10.1101/2023.06.21.545899.

51. Fleming, S.J., Chaffin, M.D., Arduini, A., Akkad, A.D., Banks, E., Marioni, J.C., Philippakis, A.A., Ellinor, P.T., and Babadi, M. (2023). Unsupervised removal of systematic background noise from droplet-based single-cell experiments using CellBender. Nat Methods 20, 1323–1335. 10.1038/s41592-023-01943-7.

52. McGinnis, C.S., Murrow, L.M., and Gartner, Z.J. (2019). DoubletFinder: Doublet Detection in Single-Cell RNA Sequencing Data Using Artificial Nearest Neighbors. Cell Syst 8, 329–337 e324. 10.1016/j.cels.2019.03.003.

53. Butler, A., Hoffman, P., Smibert, P., Papalexi, E., and Satija, R. (2018). Integrating single-cell transcriptomic data across different conditions, technologies, and species. Nat Biotechnol 36, 411–420. 10.1038/nbt.4096.

54. Emont, M.P., Jacobs, C., Essene, A.L., Pant, D., Tenen, D., Colleluori, G., Di Vincenzo, A., Jorgensen, A.M., Dashti, H., Stefek, A., et al. (2022). A single-cell atlas of human and mouse white adipose tissue. Nature 603, 926–933. 10.1038/s41586-022-04518-2.

