## Supplementary material for "CD300LG is a receptor for triglyceride-rich lipoproteins that facilitates postprandial lipid clearance": CD300LG Paper - Supplemental Figures final.pdf

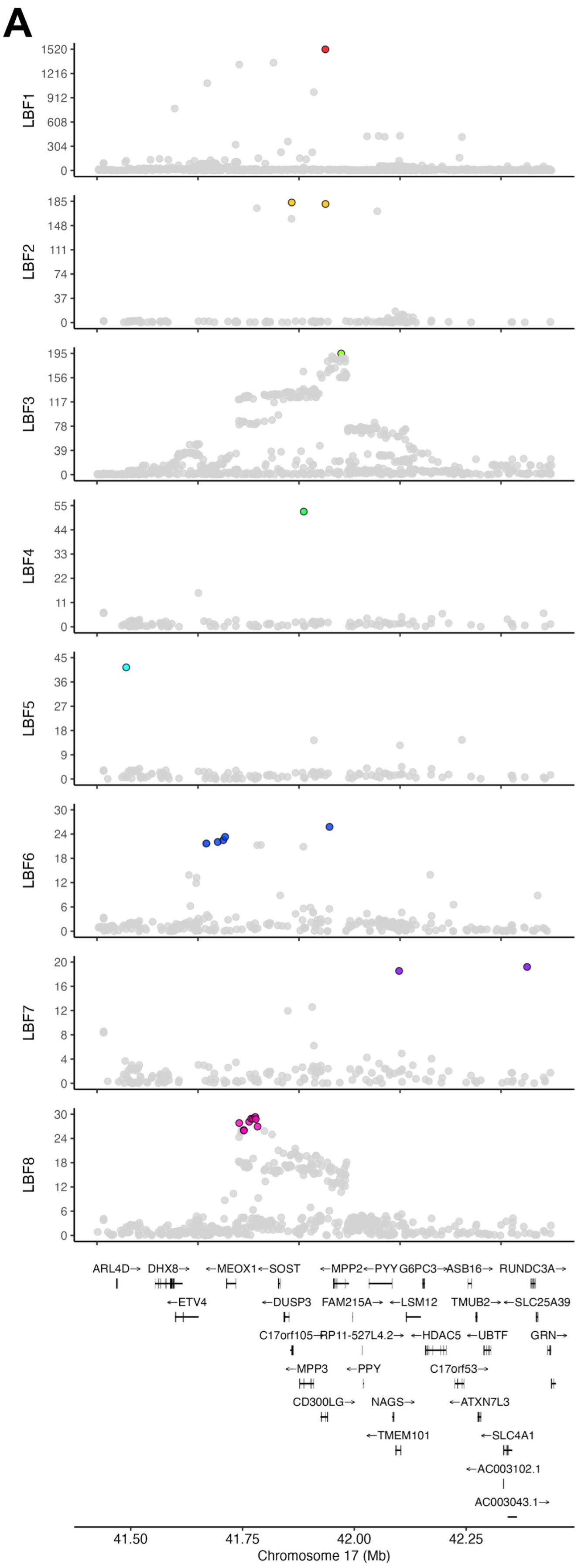


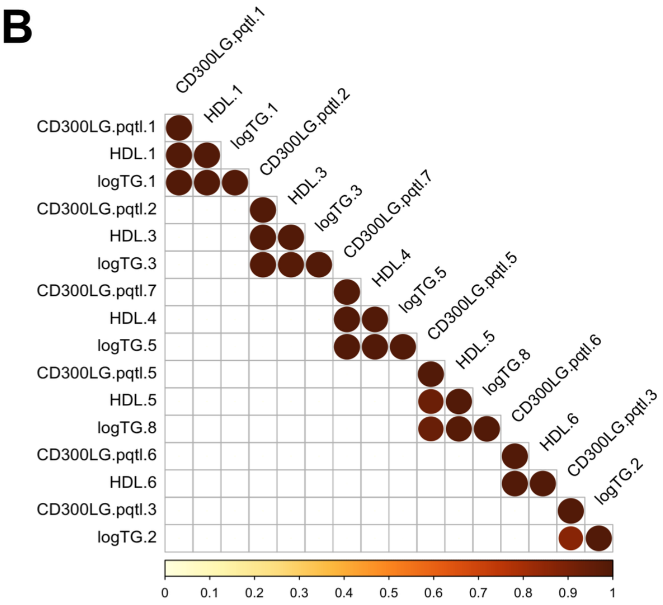


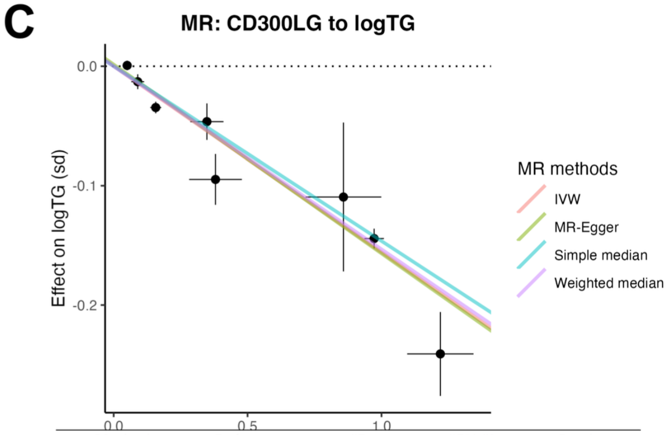


| Method | Estimate | SE | 95% CI | P-value |
|---|---|---|---|---|
| Simple median | -0.146344414 | 0.014549776 | -0.174861451 | 8.453824E-24 |
| Weighted median | -0.153258506 | 0.005505286 | -0.164048668 | 1.487249E-170 |
| IVM | -0.155723944 | 0.011055065 | -0.177391473 | 4.616966E-45 |
| MR-Egger | -0.158263752 | 0.014523928 | -0.186730127 | 0.000000E+00 |
| Intercept | 0.001485376 | 0.004910636 | -0.008139295 | 7.622852E-01 |

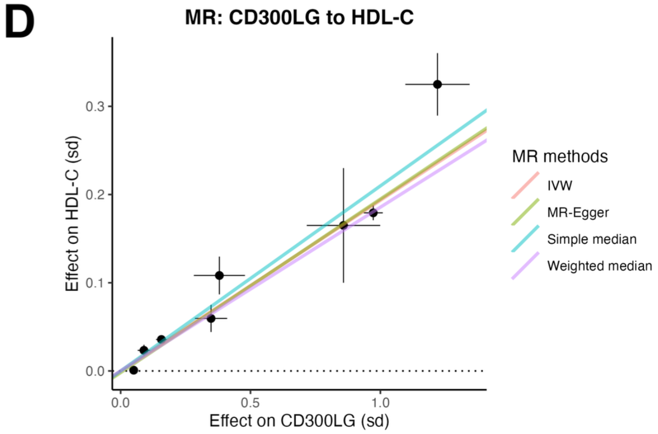


| Method | Estimate | Std Error | 95% CI | P-value |
|---|---|---|---|---|
| Simple median | 0.209756438 | 0.015620987 | 0.17913987 | 4.152361E-41 |
| Weighted median | 0.185633132 | 0.005912665 | 0.17404452 | 2.305548E-216 |
| IVM | 0.19350987 | 0.011973917 | 0.17004142 | 9.508557E-59 |
| MR-Egger | 0.196704538 | 0.015691551 | 0.16594966 | 0.000000E+00 |
| Intercept | -0.001865887 | 0.005299863 | -0.01225343 | 7.247908E-01 |

**Figure S1.** Fine-mapping, colocalization, and Mendelian randomization analyses of CD300LG cis-pQTLs. A) Fine-mapping of CD300LG GWAS summary statistics identifies 8 independent cis-pQTLs. LBF stands for log bayes factors from SuSiE fine-mapping. B) Pairwise colocalization analyses of individual CD300LG cis-pQTLs with plasma lipid signals. Color intensity represents the posterior probability of colocalization. C-D) Mendelian randomization analyses using CD300LG cis-pQTLs as exposure and triglyceride (C) or HDL cholesterol (D) levels as outcomes. The dots and error bars represent the effect size and standard error of the association between the variants and the exposure and outcome traits.

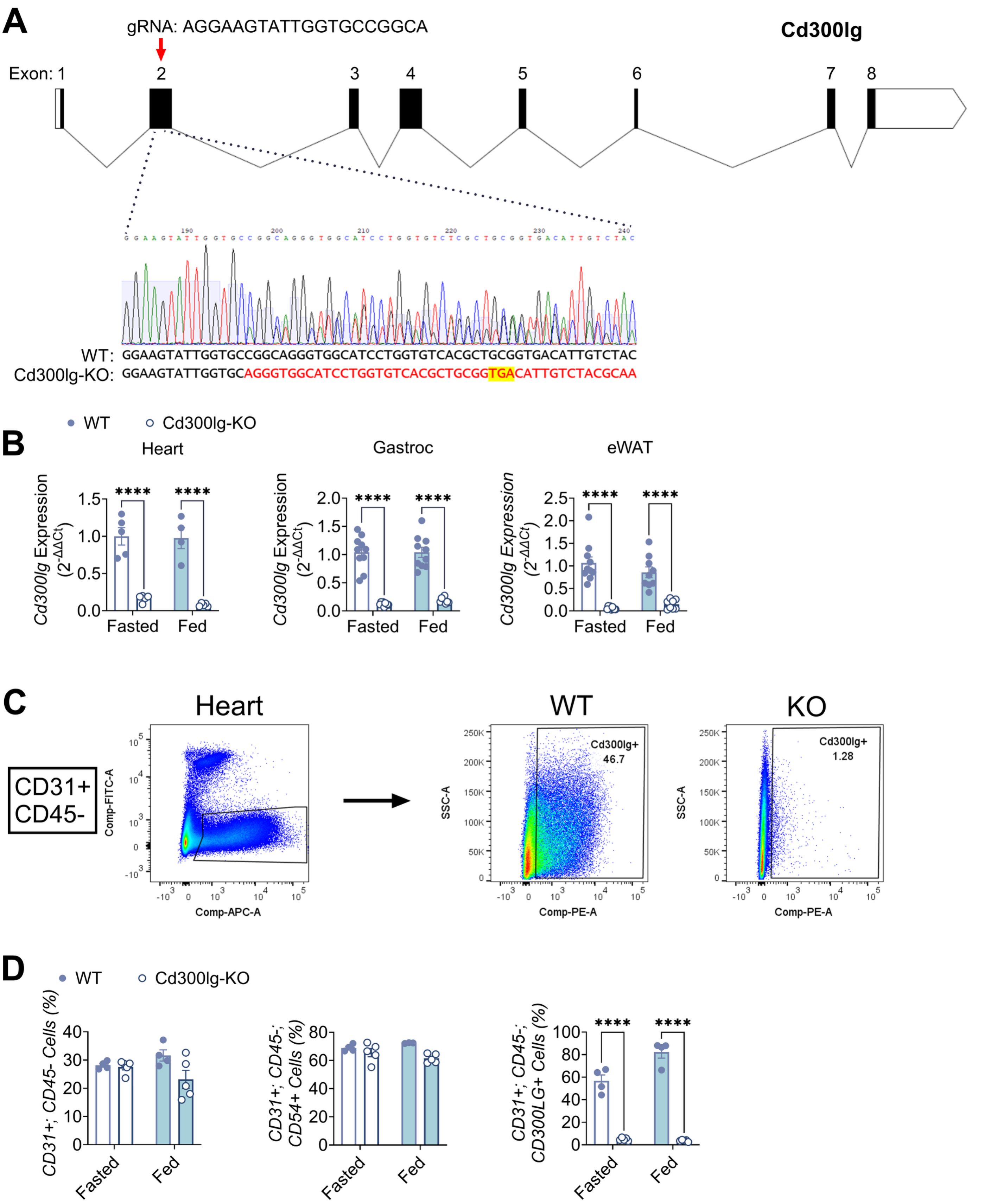


**Figure S2.** Design and validation of Cd300lg-KO mice A) Schematic of the *Cd300lg* gene, the approach and gRNAs used for knockout generation, and sequencing of the founder mouse with a 4 bp deletion (starting in red) and resulting in a premature stop codon (highlighted). B) mRNA expression of Cd300lg in WT and Cd300lg-KO mice in heart, skeletal muscle (gastroc), and epididymal adipose (eWAT), for mice that were either fasted (overnight) or fed (ad lib.). C) Scatterplots showing the gating of ECs from the heart based on CD31+ (APC) and CD45- (FITC) cells followed by gating of CD300LG+ (PE) cells in WT and Cd300lg-KO mice. D) Quantification of ECs (CD31+/CD45- or CD31+/CD45-/CD54+), and CD300LG+ ECs from WT and Cd300lg-KO mice either fasted (overnight) or fed (ad lib.). ****$p<0.0001$ by two-way ANOVA.

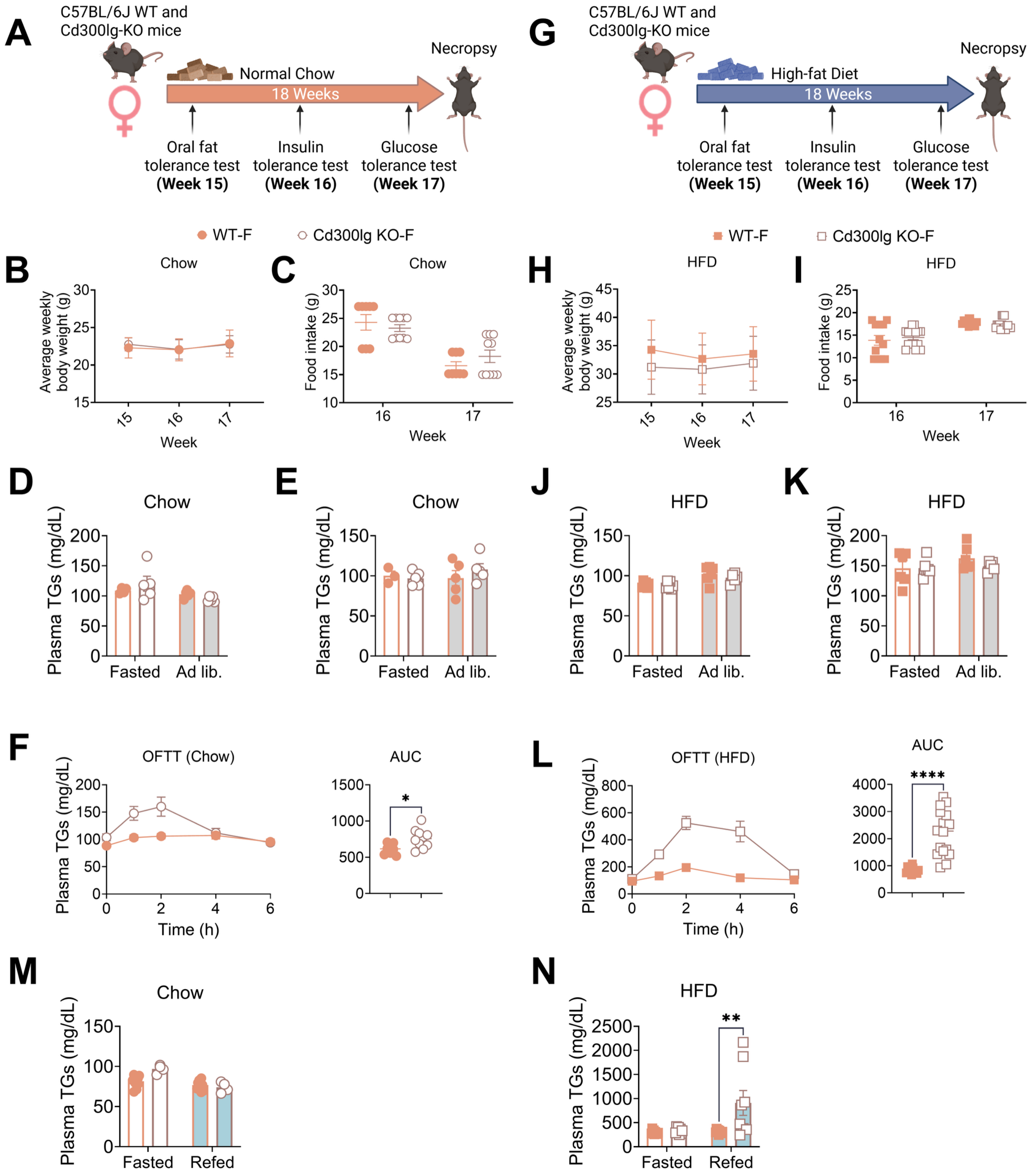


**Figure S3.** Cd300lg-KO female mice exhibit post-prandial lipid intolerance. A) Experimental design and timeline for WT and Cd300lg-KO female mice fed chow for 18 weeks post-weaning. B-C) Body weight (B) and food intake (C) for WT and Cd300lg-KO female mice on chow at weeks 15-17. D-E) Plasma TGs (D) and total cholesterol (E) in fasted (white bars) and ad libitum (gray bars) chow-fed WT and Cd300lg-KO female mice. F) Oral fat tolerance test (OFTT) in chow-fed female mice; plasma TGs were measured at the indicated time points following olive oil gavage in overnight fasted mice. G) Experimental design and timeline for WT and Cd300lg-KO female mice fed a high-fat diet (HFD) for 18 weeks post-weaning. H-I) Body weight (H) and food intake (I) for WT and Cd300lg-KO female mice on HFD at weeks 15-17. J-K) Plasma TGs (J) and total cholesterol (K) in fasted (white bars) and ad libitum (gray bars) HFD-fed WT and Cd300lg-KO female mice. L) Oral fat tolerance test (OFTT) in HFD-fed mice; plasma TGs were measured at the indicated time points following olive oil gavage in overnight fasted mice. M) Plasma TGs for WT and Cd300lg-KO female mice on chow fasted overnight or refed for 2 hours on chow. N) Plasma TGs for WT and Cd300lg-KO female mice on HFD fasted overnight or refed for 2 hours on HFD. Error bars denote S.E.M. *$p<0.05$, **$p<0.01$, ****$p<0.0001$ by t-test or two-way ANOVA as appropriate.

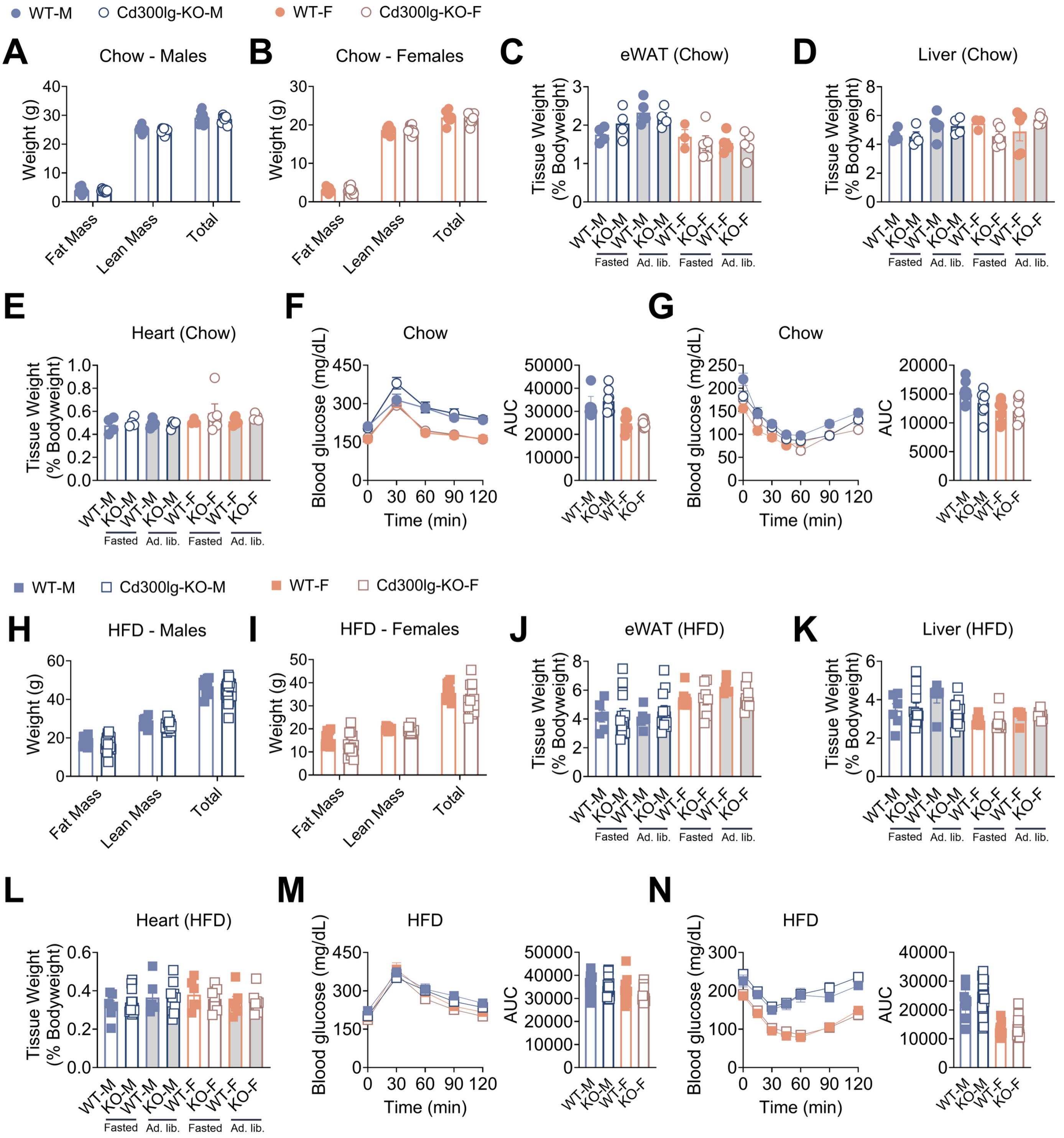


**Figure S4.** Metabolic phenotyping of Cd300lg-KO mice. A-B) Fat mass, lean mass, and total body weight by Echo-MRI for WT (closed circles) and Cd300lg-KO (open circles) male (blue)(A) and female (orange)(B) mice after 18 weeks on chow diet. C-E) Tissue weights for eWAT (C), liver (D), and heart (E), for WT and Cd300lg-KO male and female mice after 18 weeks on chow diet and either fasted overnight (white bars) or fed ad libitum (grey bars). F-G) Glucose tolerance test (F) and insulin tolerance test (G) for WT and Cd300lg-KO male and female mice on chow diet. H-I) Fat mass, lean mass, and total body weight by Echo-MRI for

WT (open squares) and Cd300lg-KO (closed squares) male (H) and female (I) mice after 18 weeks on HFD. J-L) Tissue weights for eWAT (J), liver (K), and heart (L), for WT and Cd300lg-KO male and female mice after 18 weeks on HFD and either fasted overnight or fed ad libitum. M-N) Glucose tolerance test (M) and insulin tolerance test (N) for WT and Cd300lg-KO male and female mice on HFD.

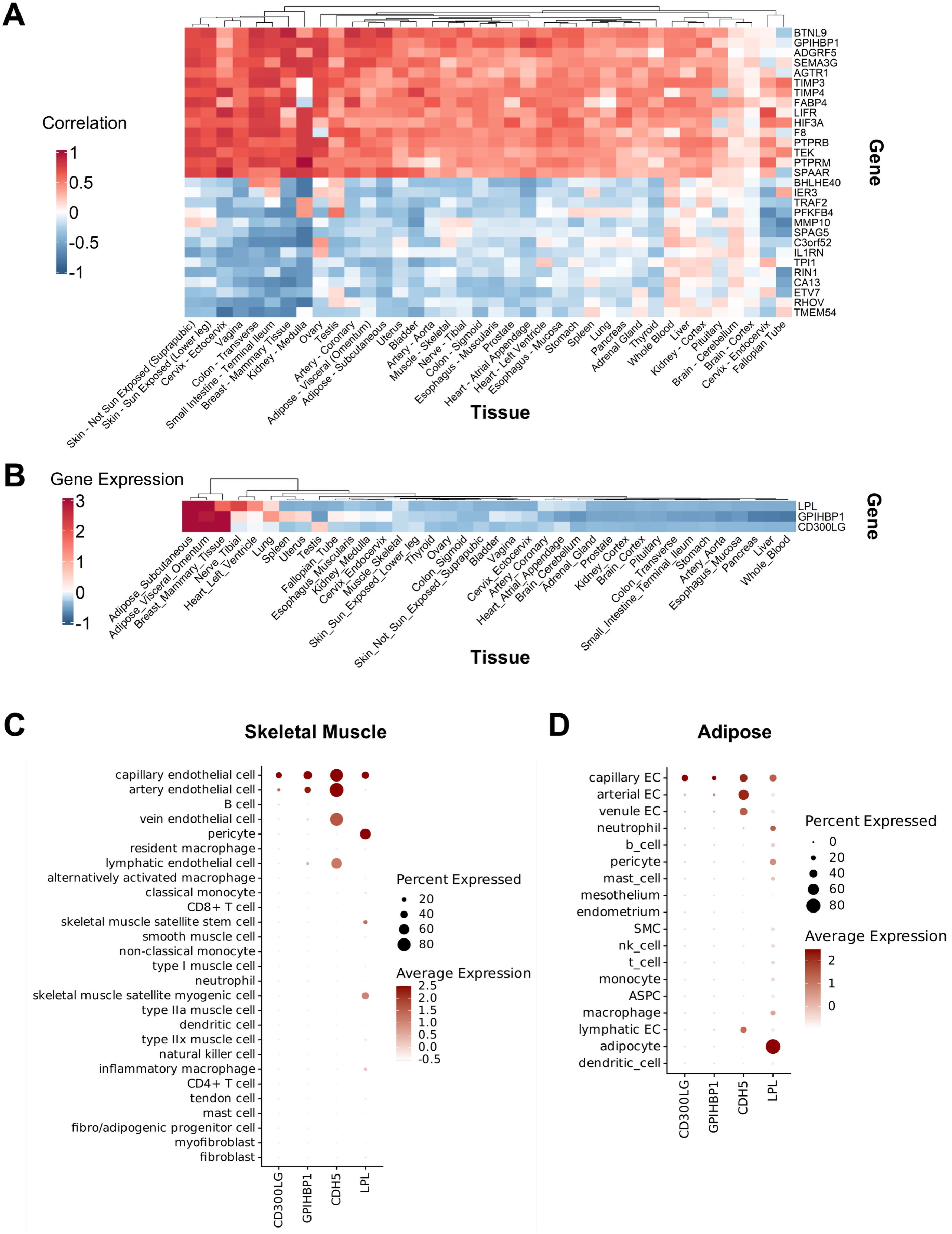

A
Correlation
1
0.5
0
-0.5
-1
BTNL9
GPIHBP1
ADGRF5
SEMA3G
AGTR1
TIMP3
TIMP4
FABP4
LIFR
HIF3A
F8
PTPRB
TEK
PTPRM
SPAAR
BHLHE40
IER3
TRAF2
PFKFB4
MMP10
SPAG5
C3orf52
IL1RN
TPI1
RIN1
CA13
ETV7
RHOV
TMEM54
Gene
Skin - Not Sun Exposed (Suprapubic)
Skin - Sun Exposed (Lower leg)
Cervix - Ectocervix
Vagina
Colon - Transverse
Small Intestine - Terminal Ileum
Breast - Mammary Tissue
Kidney - Medulla
Ovary
Testis
Artery - Coronary
Adipose - Visceral (Omentum)
Adipose - Subcutaneous
Uterus
Bladder
Artery - Aorta
Muscle - Skeletal
Nerve - Tibial
Colon - Sigmoid
Esophagus - Muscularis
Prostate
Heart - Atrial Appendage
Heart - Left Ventricle
Esophagus - Mucosa
Stomach
Spleen
Lung
Pancreas
Adrenal Gland
Thyroid
Whole Blood
Liver
Kidney - Cortex
Pituitary
Brain - Cerebellum
Brain - Cortex
Cervix - Endocervix
Fallopian Tube
Tissue
B
Gene Expression
3
2
1
0
-1
LPL
GPIHBP1
CD300LG
Gene
Adipose_Subcutaneous
Adipose_Visceral_Omentum
Breast_Mammary_Tissue
Nerve_Tibial
Heart_Left_Ventricle
Lung
Spleen
Uterus
Testis
Fallopian_Tube
Esophagus_Muscularis
Kidney_Medulla
Cervix_Endocervix
Muscle_Skeletal
Skin_Sun_Exposed_Lower_leg
Thyroid
Ovary
Colon_Sigmoid
Skin_Not_Sun_Exposed_Suprapubic
Bladder
Vagina
Cervix_Ectocervix
Artery_Coronary
Heart_Atrial_Appendage
Brain_Cerebellum
Adrenal_Gland
Prostate
Kidney_Cortex
Brain_Cortex
Pituitary
Colon_Transverse
Small_Intestine_Terminal_Ileum
Stomach
Artery_Aorta
Esophagus_Mucosa
Pancreas
Liver
Whole_Blood
Tissue
C
Skeletal Muscle
capillary endothelial cell
artery endothelial cell
B cell
vein endothelial cell
pericyte
resident macrophage
lymphatic endothelial cell
alternatively activated macrophage
classical monocyte
CD8+ T cell
skeletal muscle satellite stem cell
smooth muscle cell
non-classical monocyte
type I muscle cell
neutrophil
skeletal muscle satellite myogenic cell
type IIa muscle cell
dendritic cell
type IIx muscle cell
natural killer cell
inflammatory macrophage
CD4+ T cell
tendon cell
mast cell
fibro/adipogenic progenitor cell
myofibroblast
fibroblast
CD300LG
GPIHBP1
CDH5
LPL
Percent Expressed
20
40
60
80
Average Expression
2.5
2.0
1.5
1.0
0.5
0.0
-0.5
D
Adipose
capillary EC
arterial EC
venule EC
neutrophil
b_cell
pericyte
mast_cell
mesothelium
endometrium
SMC
nk_cell
t_cell
monocyte
ASPC
macrophage
lymphatic EC
adipocyte
dendritic_cell
CD300LG
GPIHBP1
CDH5
LPL
Percent Expressed
0
20
40
60
80
Average Expression
2
1
0

**Figure S5.** *CD300LG* gene expression correlates with *GPIHBP1* across tissue and cell types. A) Heat map of Spearman correlation values for the most positively and negatively correlated genes co-expressed with *CD300LG* using bulk tissue expression data from GTEx portal. B) Heat map of bulk tissue gene expression for *GPIHBP1*, *CD300LG* and *LPL*. C-D) Gene expression by cell type in human skeletal muscle (C) and adipose (D) for *GPIHBP1*, *CD300LG, LPL,* and the endothelial cell marker *CDH5*.

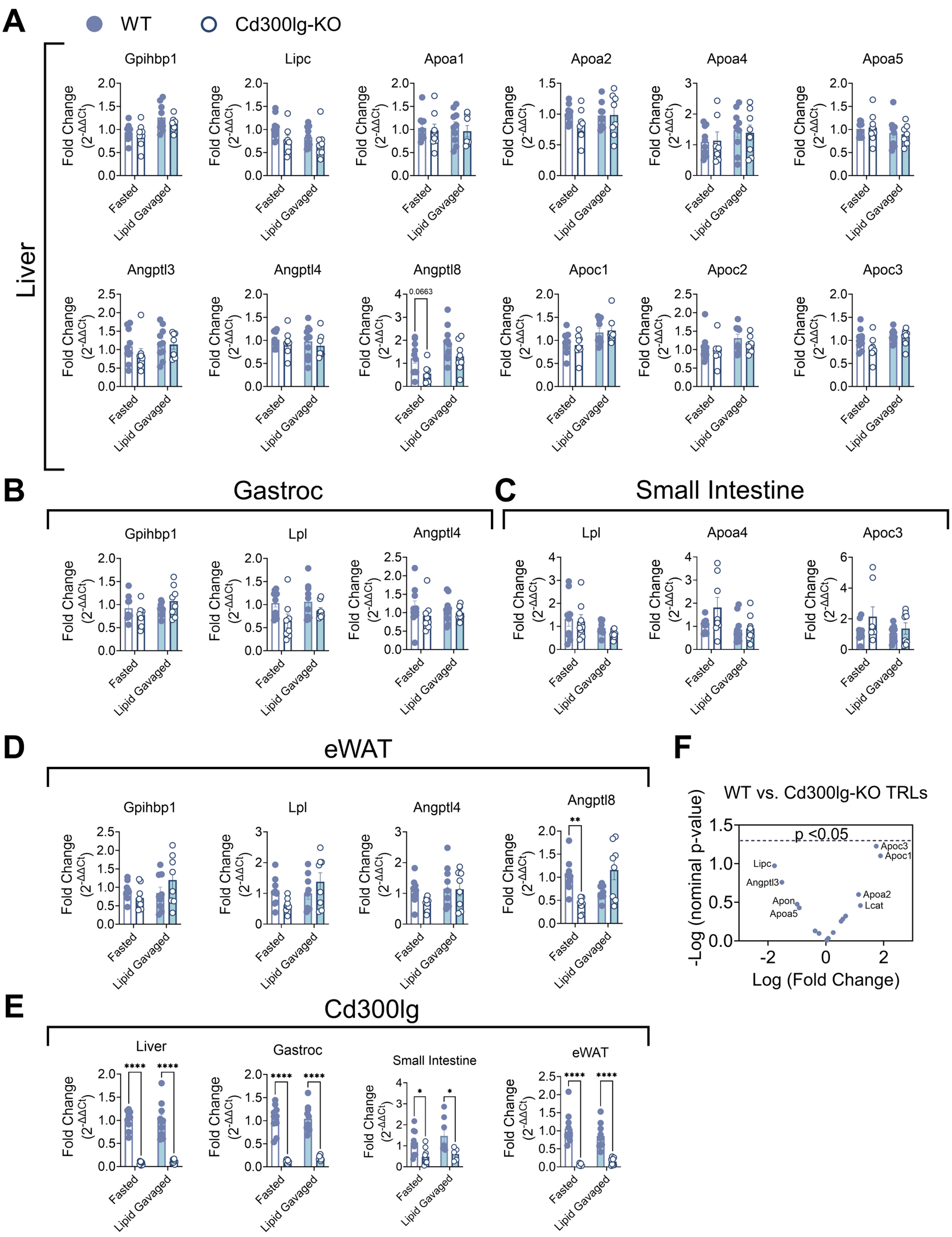
A
WT
Cd300lg-KO
Liver
Gpihbp1
Lipc
Apoa1
Apoa2
Apoa4
Apoa5
Angptl3
Angptl4
Angptl8
0.0663
Apoc1
Apoc2
Apoc3
Fold Change
($2^{-\Delta\Delta Ct}$)
Fasted
Lipid Gavaged
B
Gastroc
Gpihbp1
Lpl
Angptl4
C
Small Intestine
Lpl
Apoa4
Apoc3
D
eWAT
Gpihbp1
Lpl
Angptl4
Angptl8
**
E
Cd300lg
Liver
Gastroc
Small Intestine
eWAT
****
*
F
WT vs. Cd300lg-KO TRLs
p <0.05
Apoc3
Apoc1
Lipc
Angptl3
Apon
Apoa5
Apoa2
Lcat
-Log (nominal p-value)
Log (Fold Change)

**Figure S6.** Cd300lg-KO mice exhibit no changes in the expression of LPL regulators or TRL proteomic profile A-D) mRNA expression for known LPL regulators expressed in liver (A), skeletal muscle (B), small intestine (C), and adipose (D) of WT and Cd300lg-KO mice, following either an overnight fast (white bars) or 2 hours after an oral gavage of 10 μL/g bodyweight olive oil administered post-overnight fasting.. E) Cd300lg expression in liver, gastroc muscle, small intestine (whole), and eWAT in mice fasted overnight or 2hours after an olive oil gavage following overnight fasting. F) Proteomics analysis for TRLs isolated from WT and Cd300lg-KO mice. Dashed line denotes cutoff for $p<0.05$ adjusted by False Discovery Rate. *$p<0.05$, **$p<0.01$, ****$p<0.0001$ by two-way ANOVA.
